# EpiRanha: Hunting for Epitope Similarity with a Structure- and Residue-Aware Graph Neural Network

**DOI:** 10.64898/2026.04.21.719830

**Authors:** Tygo Francissen, Miriam Babukhian, Helena Britze, Yano Wilke, Roberto Spreafico, Samuel Demharter, Marloes Arts

**Author notes:** These authors contributed equally (Tygo Francissen, Miriam Babukhian).

## Abstract

Precise epitope recognition underpins the efficacy and safety of therapeutic antibodies, yet existing approaches to epitope similarity scoring rely largely on sequence identity or rigid structural superposition, limiting their ability to robustly assess cross-reactivity and potential off-target interactions. We introduce **EpiRanha**, a multimodal framework that integrates residue-level ESM-2 sequence embeddings with an E(n)-equivariant graph neural network operating on three-dimensional protein structure. EpiRanha produces per-residue “fingerprints” that jointly encode sequence-level context and spatial organization, and then applies a beam-search strategy to identify and rank multiple high-confidence epitope candidates across protein surfaces that are similar to a given query epitope. We evaluate EpiRanha against TM-align on nanobody-antigen complexes from SAbDab-nano and a set of AlphaFold-predicted proteins. EpiRanha consistently recovers the query epitope on its cognate antigen, including highly discontiguous conformational epitopes that rigid alignment methods such as TM-align often fail to capture, achieving lower structural loss and fewer false negatives through flexible residue-level mapping. Beyond these self-matches, EpiRanha also identifies biologically plausible epitope-level similarities on other proteins. Overall, EpiRanha advances epitope characterization beyond sequence or geometry alone, enabling more robust off-target risk assessment, informing training-set construction for predictive models, and more selective antibody design.

## 1 Introduction

The success of a therapeutic antibody requires both high target specificity and minimal cross-reactivity [1, 2, 3]. In this context, the target is a specific epitope, i.e., a particular binding region on a given antigen. Accurate computational characterization and quantitative similarity scoring of epitopes are critical for advancing vaccine and therapeutic antibody design, improving target selectivity, and identifying epitope-like sites in unrelated proteins that may cause off-target effects [4, 5, 6, 7]. *Epitope matching*, which we define as the task of searching for similar epitopes across a protein database (see Figure 1a), is instrumental for predicting potential cross-reactivity and identifying conserved recognition sites [5, 6, 7].

**Figure 1:**
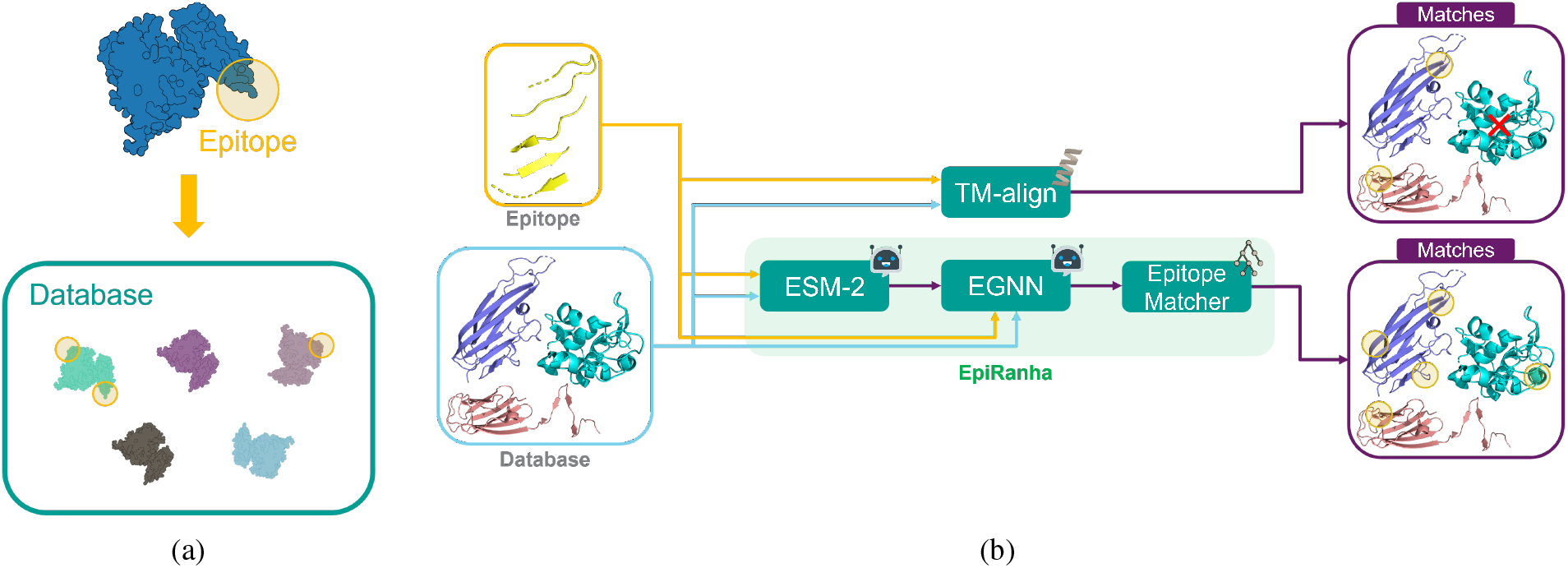
Overview of the epitope matching problem and proposed methods. (a) “Epitope matching” is the task of detecting structurally and functionally similar epitope regions (highlighted in yellow) in proteins across a database. (b) Comparison of epitope matching approaches: TM-align performs rigid backbone alignment, while EpiRanha integrates sequence embeddings (ESM-2), geometric message passing (EGNN), and beam search to predict flexible, learned epitope similarity. Robot icons represent ML-based components.

Most existing similarity metrics are primarily based on sequence identity [8, 9]. However, such approaches often fail to account for the fact that distinct primary sequences can fold into highly similar, or even identical, 3D epitopes. This phenomenon is well-supported by the success of inverse folding models [10, 11], which demonstrate that a single target fold can be successfully stabilized by a vast and diverse array of amino acid sequences. The effectiveness of epitope matching is therefore significantly reduced for purely sequence-based methods, which also overlook the structural and chemical determinants that govern molecular recognition. This poses a particular challenge for so-called *conformational epitopes*, which, as opposed to linear epitopes, comprise discontinuous residues brought into proximity by protein folding [12].

Antibody-antigen binding is highly dependent on three-dimensional structure: the interaction is dictated by a distributed network of weak interactions, including hydrogen bonds, van der Waals contacts, and electrostatic forces, across a chemically diverse and solvent-exposed interface [13, 14]. As a result, the computational characterization of conformational epitopes depends on an accurate three-dimensional representation of the antigen surface and its structural context [4, 13].

Structural alignment tools like TM-align [15] have become instrumental in protein structure comparison, facilitating the identification of conserved structural features critical for functional inference, protein engineering, and drug design [16, 17]. TM-align is a rigid-body algorithm that identifies optimal alignments through a single global rotation and translation, aiming to maximize structural overlap based solely on protein backbone geometry [18, 19]. However, this rigid-body approach has inherent limitations, producing only a single global alignment and remaining computationally demanding for large-scale analyses. In contrast, high-throughput structural motif search frameworks such as Folddisco [20] enable rapid scanning of millions of structures and are specifically designed to detect spatially conserved residue patterns, including discontinuous segments (i.e., structural discontiguity) that lack linear sequence correspondence. Such approaches illustrate the advantages of scalable, motif-centric methods over traditional global alignment strategies. Nevertheless, even motif-level structural searches remain predominantly geometry-driven and may overlook physicochemical and evolutionary determinants of functional similarity. To support robust predictive modeling and the safe, reliable design of molecular binders, epitope-similarity metrics must therefore integrate high-resolution 3D structural information with residue-level contextual descriptors that reflect sequence and evolutionary constraints [4, 5, 14, 21].

To overcome the limitations of methods relying exclusively on protein sequence or atomic structure, recent machine learning approaches have introduced diverse representations that capture complementary aspects of protein biology. Methods such as MaSIF and dMaSIF [22, 23] apply geometric deep learning to capture molecular surface features, enabling the analysis of shape complementarity and physicochemical properties at the protein interface. In parallel, large-language models (LLMs) like ESM-2 [24] are able to learn both evolutionary and biochemical information from protein sequences alone. While these approaches provide powerful new ways of representing proteins, they also reflect distinct modeling emphases: methods such as MaSIF focus on geometric and physicochemical features of protein surfaces, whereas sequence-based models such as ESM-2 primarily encode residue-level evolutionary and biochemical information without explicit spatial representation.

In an effort to bridge this divide, several approaches integrate structural and sequence information. Many were developed for general protein-protein interactions (PPIs), while others, such as GraphBepi [25], focus specifically on epitope prediction by combining AlphaFold-derived structures with ESM-2 embeddings. Despite these advances, general PPI models are not optimized for antibody-antigen recognition, and epitope-focused models often emphasize predictive labeling rather than flexible, residue-level matching across multiple antigens. A more detailed overview of related work and methodological comparisons is provided in Appendix A.

Building on these insights, we introduce EpiRanha (“Epi” refers to epitopes, while “(pi)Ranha” evokes rapid and precise target identification): a valuable new tool in the epitope matching toolbox that overcomes the limitations of methods like TM-align. An overview is shown in Figure 1b. EpiRanha integrates three-dimensional structural information with ESM-2 embeddings, which capture amino acid sequence information and implicit physicochemical features. By combining these modalities within a single representation, it learns residue-level fingerprints that reflect both spatial organization and biochemical context. On top of this, a beam-search-based strategy enables the identification of not just a single alignment but multiple candidate matches for a given query epitope, ranking them according to their structural and embedding similarity and overcoming the single-solution limitation of rigid aligners. This flexible approach provides a more robust basis for epitope similarity scoring than methods relying on sequence or structure alone.

With this work, we establish a robust framework for epitope characterization and similarity scoring, with direct implications for therapeutic nanobody and antibody design. Our contributions include enabling the assessment of off-target cross-reactivity and advancing the broader understanding of epitope features and key properties of molecular binding. These tools can also guide the construction of diverse training sets to support downstream predictive tasks in machine learning models. Moreover, we present a novel hybrid sequence-structure model for protein similarity scoring that captures complementary determinants of molecular recognition.

## 2 Methods

### 2.1 Data Collection

The primary component of our dataset consists of nanobody-antigen pairs sourced from SAbDab-nano [26], a publicly available database containing all nanobody-antigen complexes available in the Protein Data Bank (PDB) [27], providing both sequence and experimentally resolved structural information. For representation learning only, we supplemented the dataset with AlphaFold 2-predicted surface proteins [28]. Detailed descriptions of the dataset construction are provided in Appendix B. Basic descriptive statistics of epitope and antigen lengths are shown in Supplementary Appendix M.1.

### 2.2 TM-align

We benchmarked EpiRanha against TM-align [18], a widely used backbone-based structural alignment method. TM-align was applied to all epitope-antigen pairs in an all-vs-all manner, aligning each epitope structure against every antigen structure in the benchmark set using only *C*_*α*_ coordinates. All alignments were performed using TM-align (version 20220227) with default parameters. No additional preprocessing or residue filtering was applied prior to alignment beyond the PDB structures provided as input. TM-align was used solely to generate backbone-based structural alignments and residue-level correspondences. For each epitope–antigen pair, only the residues identified as aligned by TM-align were retained and used for subsequent evaluation, as described in Section 2.4.

### 2.3 EpiRanha

EpiRanha operates in two main stages: fingerprinting and epitope matching. In the first stage, protein fingerprints are created by combining structural and sequence-derived information. In the second stage, these fingerprints are used to identify structurally and functionally similar regions across proteins. Figure 2 provides an overview of these two stages.

**Figure 2:**
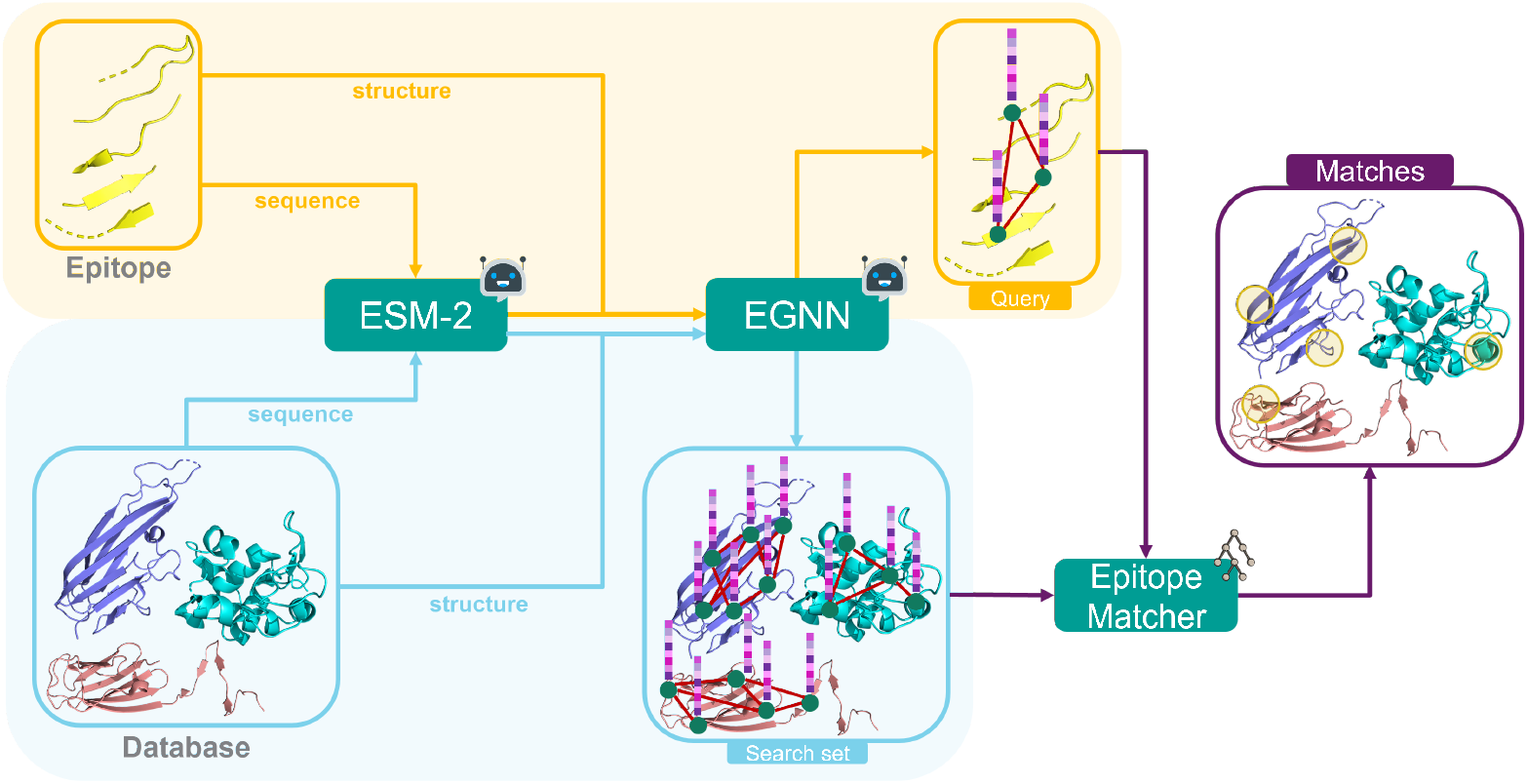
Overview of the EpiRanha workflow. In the fingerprinting stage, both the query epitope and a set of search proteins are represented through two complementary modalities: sequence information encoded by fixed, pretrained ESM-2 embeddings, and structural information encoded by an equivariant graph neural network (EGNN). The ESM-2 embeddings are propagated through the EGNN to yield per-residue fingerprints that integrate sequence and spatial context. In the matching stage, the query and search-set fingerprints are compared in the epitope matcher, which identifies structurally and functionally similar regions using a beam search strategy. The highest-scoring matches correspond to candidate epitopes with similar geometric and biochemical properties. Robot icons represent ML-based components.

#### 2.3.1 Fingerprinting

The fingerprinting stage of EpiRanha employs a multimodal approach that combines a graph neural network with ESM-2 embeddings [29]. Specifically, an equivariant graph neural network (EGNN) [30] is employed to preserve equivariance to translations, rotations, reflections, and permutations of input atomic coordinates, ensuring that structural relationships remain consistent across transformations. Inspired by methods such as MaSIF [22], our EGNN is designed to produce low-dimensional, per-residue embeddings that retain both structural and sequential information.

The protein is represented as a residue-level graph, where nodes are placed at residue C_*α*_ atomic coordinates. Edges are defined based on spatial proximity, controlled by a tunable parameter that determines the number of nearest neighbors per residue. Each node is enriched with ESM-2 embeddings, providing a powerful sequence-based descriptor learned from large-scale protein language models.

To train the EGNN, we implement a masking strategy where a certain number of residues and their corresponding ESM-2 embeddings are randomly masked out during training. Further details are provided in Appendix C.1 and Figure C1. The model has to reconstruct these missing nodes and their features through message passing, forcing it to learn both the contextual relationships between residues within a protein and the sequential and structural dependencies between nodes.

Training is performed using a composite loss function consisting of a coordinate reconstruction loss and a feature reconstruction loss. The coordinate loss is defined as the mean squared error between predicted and true residue coordinates, while the feature loss is defined as the mean squared error between predicted and original ESM-2 embeddings. The total loss is a weighted sum of these two terms, with weights chosen to ensure balanced contributions during optimization, as further detailed in Appendix C.2. Once the model is trained to convergence, the per-residue features are extracted from the EGNN, just before the final layer. Since they have been updated based on spatial interactions with neighboring nodes, they integrate both sequential and structural information. More details and a complete overview of the EGNN training workflow, from raw protein input to per-residue embedding generation, is provided in Appendix C.

To verify stable training and meaningful representation learning, we evaluated EGNN reconstruction behavior using both quantitative metrics and qualitative visualizations. These reconstruction-based sanity checks assess whether the model consistently captures local geometric and sequence context and are not interpreted as performance metrics for epitope matching. Overall, the reconstruction results indicate stable training and that the model reliably captures local geometric and sequence context. The corresponding analyses are reported in Appendix D.

#### 2.3.2 Epitope Matching

In the second stage, EpiRanha uses these embeddings to identify regions in other proteins (“search”) that resemble a given epitope (“query”). Conceptually similar to deploying TM-align in a one-versus-many setup, this stage scans a query epitope against a search set of proteins to detect structurally and functionally analogous regions.

For each search protein, all per-residue embeddings and coordinates are precomputed and cached. EGNN embeddings used for epitope matching were trained with a masking fraction of 0.1 on a combination of SAbDab-nano antigens and AlphaFold-predicted proteins. Beam search parameters, including the number of nearest neighbors per residue and the beam width, were chosen based on qualitative benchmarking on representative proteins to balance computational efficiency with robust epitope detection. Appendix E provides detailed analyses of parameter sensitivity, ablation studies, and sanity checks that motivated these choices, including exploration of masking fractions, beam widths, and neighbor counts across different query-search protein sizes.

Within EpiRanha, epitope matching is formulated as a path-finding problem over the residue graph of the search protein, where each path corresponds to a candidate epitope-sized region. To efficiently explore this large search space, EpiRanha employs beam search. The algorithm initializes a beam from every residue in the search protein, making the procedure inherently parallelizable. From each starting point, the algorithm incrementally expands candidate paths by adding spatially adjacent residues, where neighborhood relationships are defined by a tunable number of nearest neighbors in 3D space.

Each partial or complete path is evaluated using a combined similarity function that considers both residue-level features and structural correspondence. Given a candidate path and query epitope, an optimal one-to-one residue mapping is first computed by solving a linear sum assignment problem using a modified Jonker-Volgenant algorithm without initialization [31]. Based on this mapping, two distance-based terms are calculated: a structural term, which measures the Euclidean distances between matched residue coordinates, and a feature term, which measures the Euclidean distances between the corresponding residue embeddings. The total score of a candidate path is defined as a weighted sum of these two terms, with weights calibrated by first evaluating each term independently and then scaling them to ensure comparable contributions. Formally, this scoring function corresponds to the assignment-based similarity score defined in Appendix F.1.

During beam search, only the top-scoring candidate paths (as defined by the beam width) are retained for further expansion, while lower-scoring paths are pruned early. This iterative pruning ensures that the search remains tractable without compromising accuracy.

Path extension continues until a path reaches the same length as the query epitope, at which point it is considered a complete match. Final matches are ranked according to their similarity scores, and the highest-scoring regions are selected for downstream analysis. A visual overview of the epitope matching workflow and detailed pseudocode are provided in Appendix F.

### 2.4 Evaluation

Because ground truth data indicating correct cross-protein epitope matches are scarce, evaluation is performed on individual epitope-antigen and epitope-protein pairs. Two main categories are considered: 1) *self-matching* (or cognate matching), where epitopes are aligned against their original antigens, serving as a baseline where perfect matches are expected, and 2) *cross-matching*, where epitopes are aligned against other proteins to assess how different methods identify structurally and biochemically similar surface regions.

Importantly, evaluation is decoupled from the search procedure itself. Both TM-align and EpiRanha first produce candidate residue sets (paths) on the search protein. These candidate paths are then evaluated using a unified, post hoc scoring procedure to assess how well the identified surface patch matches the query epitope.

#### 2.4.1 Path-Based Evaluation and Residue Mapping

Both EpiRanha and TM-align ultimately aim to identify a surface patch on a search protein that corresponds to a given query epitope. To enable a fair comparison, evaluation is performed solely on the set of residues selected by each method, independent of how that set was obtained.

For EpiRanha, each candidate match is represented as a path of residues produced by beam search. For TM-align, aligned residue positions are extracted from the structural alignment output and treated as candidate paths. In both cases, only the residue indices are retained; no fixed residue-to-residue correspondence is assumed at this stage. This design choice reflects the primary objective of the task: identifying which region of the protein surface corresponds to the epitope, rather than enforcing a particular alignment during search.

To assess match quality, an optimal one-to-one mapping between the query epitope residues and the candidate path residues is computed after candidate generation. This mapping is obtained by matching residues between the query and the candidate according to their structural and feature-level similarity. Importantly, this mapping step is not part of the search procedure itself but is applied uniformly during evaluation, ensuring that match quality reflects the intrinsic similarity between the two residue sets rather than the specifics of the search strategy.

#### 2.4.2 Match Quality Metrics

Final match quality is assessed using two complementary metrics, computed under the optimal residue mapping described above.

Coordinate loss quantifies structural similarity by summing the Euclidean distances between the *C*_*α*_ coordinates of matched residue pairs. Lower coordinate loss indicates stronger geometric agreement between the identified surface patch and the query epitope.

Feature-based loss is used to assess residue-level similarity that is not captured by geometric alignment alone. Because embedding-based feature spaces differ in scale and geometry, feature losses are not compared directly across different embedding types. Instead, BLOSUM-based evaluation provides an embedding-independent reference grounded in observed amino-acid substitution patterns. For that, substitution scores are obtained from the BLOSUM62 matrix. Raw substitution scores are converted into cost values by subtracting each score from the maximum attainable BLOSUM score, such that lower costs correspond to higher biochemical similarity. Under this formulation, even strong matches incur a nonzero cost, and total BLOSUM-based losses are therefore expected to remain positive.

The total evaluation score is defined as a weighted combination of coordinate loss and feature-based loss, where higher scores indicate better matches. Formally, the evaluation computes the negative of the assignment-based similarity score defined in Appendix F.1, replacing the beam search feature embeddings with evaluation-specific features. In particular, BLOSUM-based evaluation replaces embedding distances with BLOSUM-derived substitution costs and uses a corresponding set of fixed coordinate and feature weights. All weights are fixed a priori, applied uniformly across methods, and listed in Table F1. These metrics are not optimized during model training or beam search and therefore serve as independent measures of match quality.

To calculate a per-match confidence score, a set of five random decoy matches per query-search pair was generated together with their corresponding coordinate losses. These losses were pooled and used as a reference, such that for each match we can provide a *p*-value and a corresponding confidence score −log_10_(*p*) that indicates the chance of randomly finding a match with the same loss. The *p*-value is defined as:

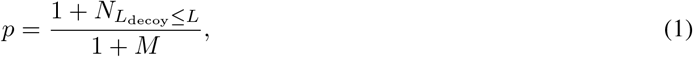

where 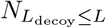 is the number of decoys with a loss equal or lower to the current match loss *L*, and *M* is the total number of decoys.

#### 2.4.3 Loss Normalization

To enable fair comparison across methods and across matches of varying epitope length, all alignment-based losses were normalized after computation. This normalization accounts for both alignment coverage and loss scale and is applied consistently across methods and loss types. For TM-align, only residues included in the structural alignment contribute to the reported alignment and associated loss values, consistent with how TM-align computes its alignment statistics. In low-confidence cases, this may result in short alignments that cover only a subset of the query epitope, which can lead to artificially favorable scores when losses are computed over the full chain length. In contrast, EpiRanha produces full-length epitope paths and therefore inherently accounts for all residues. To mitigate this discrepancy, we introduce a full-length–normalized loss formulation that explicitly penalizes unmatched residues. Let *L* denote the full epitope length, *a* the number of residues included in the alignment, and *S* the cumulative loss over aligned residues. For EpiRanha, *a* = *L*, while for TM-align, *a* corresponds to the aligned length reported by the method. The normalized loss is defined as:

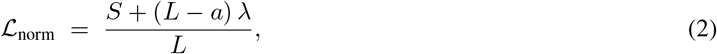

where *λ* is a fixed per-residue penalty applied to each unmatched epitope residue (see Appendix G). This formulation yields a per-residue loss that jointly reflects alignment quality and epitope coverage. Under this formulation, unmatched residues are penalized implicitly through the coverage term (*L* − *a*). For EpiRanha, the aligned length equals the full epitope length (*a* = *L*), and therefore no unmatched-residue penalty is incurred. For the BLOSUM-based loss, raw BLOSUM62 substitution scores were first shifted by subtracting the minimum achievable loss for each epitope–antigen pair under the corresponding residue mapping, such that the best possible match for that pair corresponds to zero loss prior to normalization. This ensures that all normalized losses are comparable across epitopes with different residue compositions.

#### 2.4.4 Epitope Discontiguity

Epitope discontiguity refers to an epitope composed of physically separated regions of the antigen, resulting in a fragment that is not continuous in either the primary sequence or the three-dimensional structure once isolated. We quantified discontiguity by computing a raw score *D* that accounts for both the number and sizes of any gaps, normalized by the total epitope length. Specifically, let *N* be the number of gaps, *s*_*i*_ the size of gap *i, α* and *β* tunable weights, and *L* the epitope length. We define

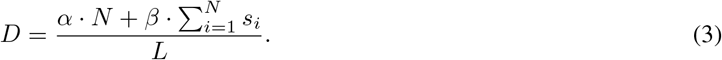

To limit this raw score between 0 and 1, we apply a rational saturating (hyperbolic) transformation of the form:

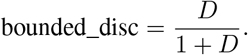

Hence, higher values of *D* yield bounded discontiguity scores closer to 1, indicating greater structural fragmentation. In this work, we set *α* = 2 and *β* = 1, thereby penalizing the introduction of new gaps more strongly than the extension of existing gaps. This choice is conceptually motivated by affine gap penalty models commonly used in sequence alignment, where opening a gap incurs a larger penalty than extending it [32]. While our discontiguity score is not an alignment objective, this weighting reflects the intuition that multiple separate fragments indicate greater epitope fragmentation than a single extended gap of comparable total length (see Supplementary Appendix M.2).

### 2.5 Experimental Setup

Computational infrastructure and execution environment details are described in Appendix H.

## 3 Results

### 3.1 Comparative Epitope Matching Performance

We compare EpiRanha to TM-align on the task of epitope matching using a benchmark of experimentally resolved epitopes and their corresponding antigens. Each epitope was aligned against every antigen in an all-vs-all setting, yielding a dense set of epitope-antigen comparisons. Performance was evaluated using two complementary metrics: a coordinate-based loss measuring geometric agreement between matched residues, and a BLOSUM-based loss capturing evolutionary plausibility under an optimal residue mapping.

The resulting similarity heatmaps summarize these losses for all epitope-antigen pairs. Diagonal entries correspond to self-matching cases, where each epitope is aligned to its native antigen, while off-diagonal entries reflect cross-antigen similarity and potential non-cognate matches. Because epitopes are defined from experimentally resolved bound structures, based on antibody occupancy on the antigen surface, a few antigens contribute multiple distinct epitopes, resulting in 146 epitopes derived from 142 corresponding antigens.

#### 3.1.1 Self-Matching Performance

Figures 3 and 4 show the coordinate-based and BLOSUM-based loss heatmaps for TM-align and EpiRanha, respectively. TM-align was designed to optimize structural alignment and does not incorporate residue-level features, whereas EpiRanha leverages both geometry and biochemical information. To capture these complementary aspects, we report two metrics: coordinate loss, measuring the mean squared error between predicted and true residue coordinates, and BLOSUM feature loss, assessing amino-acid substitution plausibility.

**Figure 3:**
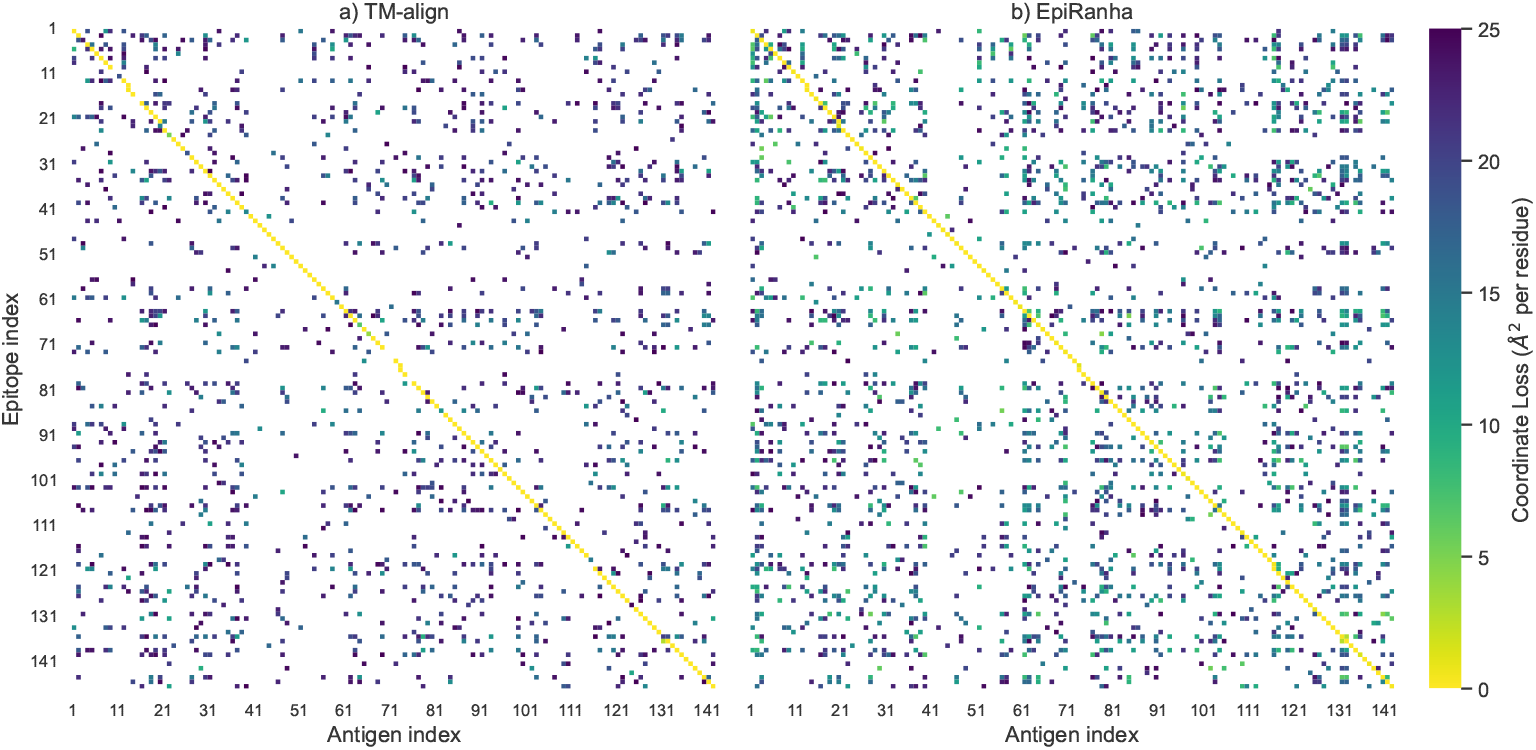
Structural alignment loss for TM-align and EpiRanha across antigens and epitopes. Heatmaps show per-epitope coordinate-based loss for 146 epitopes (rows) aligned against 142 antigens (columns) using (a) TM-align and (b) EpiRanha. The diagonal corresponds to self-matching cases, where each epitope is aligned to its native antigen, while off-diagonal entries represent alignments to non-cognate antigens. Residue-wise coordinate loss was capped at 25 Å^2^ for visualization. Lower values indicate better structural agreement. For readability, only every *k*-th epitope and antigen index is labeled; complete mappings between heatmap indices and PDB identifiers (including antigen lengths) are provided in Appendix M.3.

**Figure 4:**
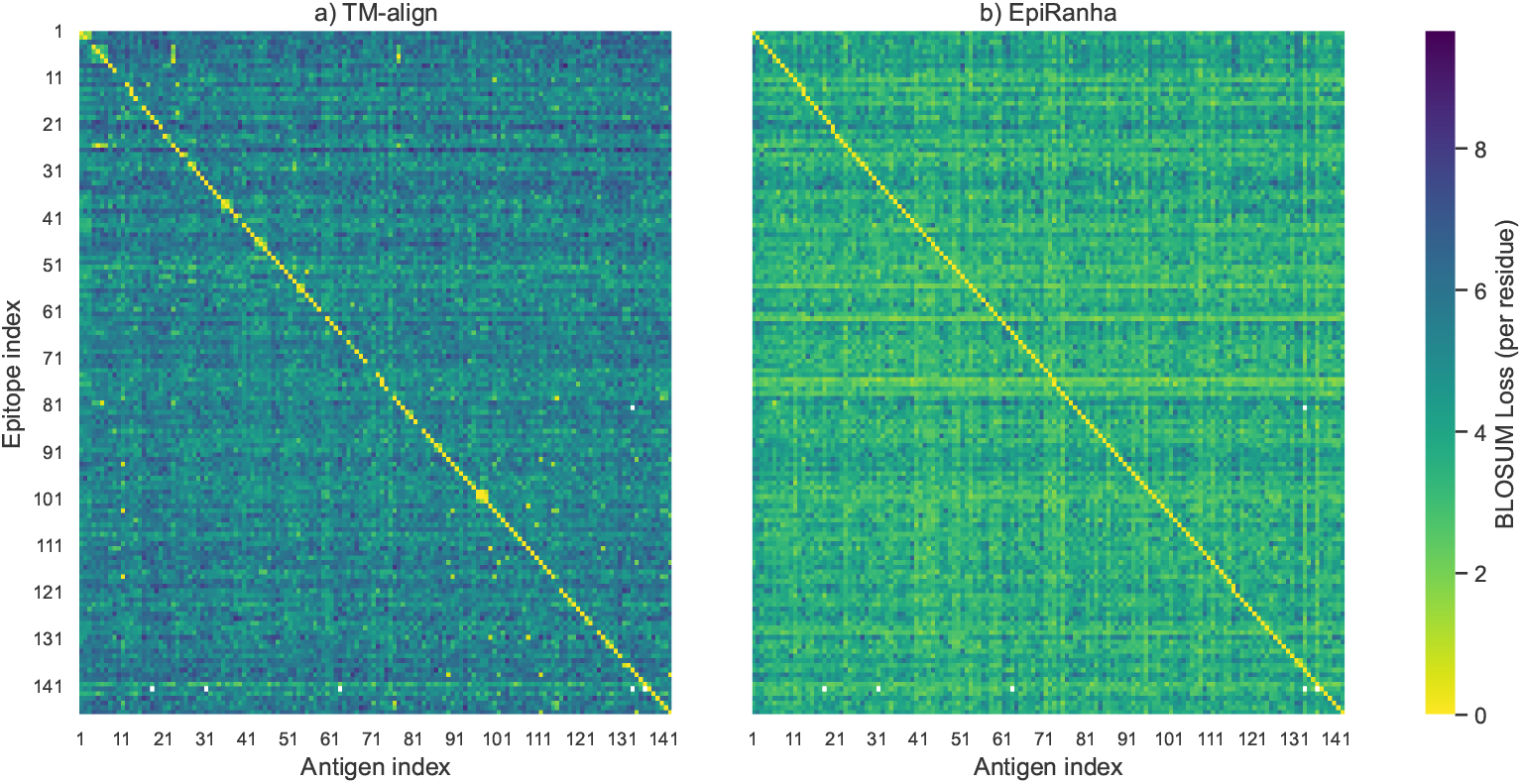
BLOSUM-based feature alignment loss for TM-align and EpiRanha across antigens and epitopes. Heatmaps show per-epitope BLOSUM loss for the same epitope–antigen pairs as in Figure 3, reflecting biochemical similarity under an optimal residue mapping. The diagonal indicates self-matching cases, while off-diagonal entries correspond to cross-antigen alignments. Lower values indicate higher biochemical compatibility. Axis labeling and index–PDB mappings are as in Figure 3 (see Appendix M.3).

For the coordinate-based metric, since coordinate discrepancies are evaluated using a mean squared error, large outlier distances can disproportionately dominate the loss. We therefore cap pairwise residue distances at (5^2^ = 25) Å^2^, corresponding to the squared length scale of a generous single-residue positional error. Given that the characteristic spatial extent of an amino acid residue is on the order of a few angstroms, this threshold allows for substantial local flexibility while preventing implausible long-range mismatches from overwhelming the loss.

In both methods, the strongest signal appears along the diagonal, reflecting the recovery of each epitope on its native antigen. EpiRanha consistently exhibits near-zero loss for these pairs in both structure- and feature-based heatmaps, indicating robust self-matching performance. TM-align also recovers many self-matches, but shows clear gaps and elevated losses, reflecting missed or partial alignments even in this favorable setting.

These differences are quantified in Figure 5a. EpiRanha achieves near-zero loss for almost all epitopes, while TMalign fails to recover approximately 20 cognate matches, resulting in a substantial tail of high-loss outliers with per-residue deviations ranging from roughly 5 to 40 Å^2^. These failures correspond directly to the gaps observed on the diagonal of the heatmaps.

**Figure 5:**
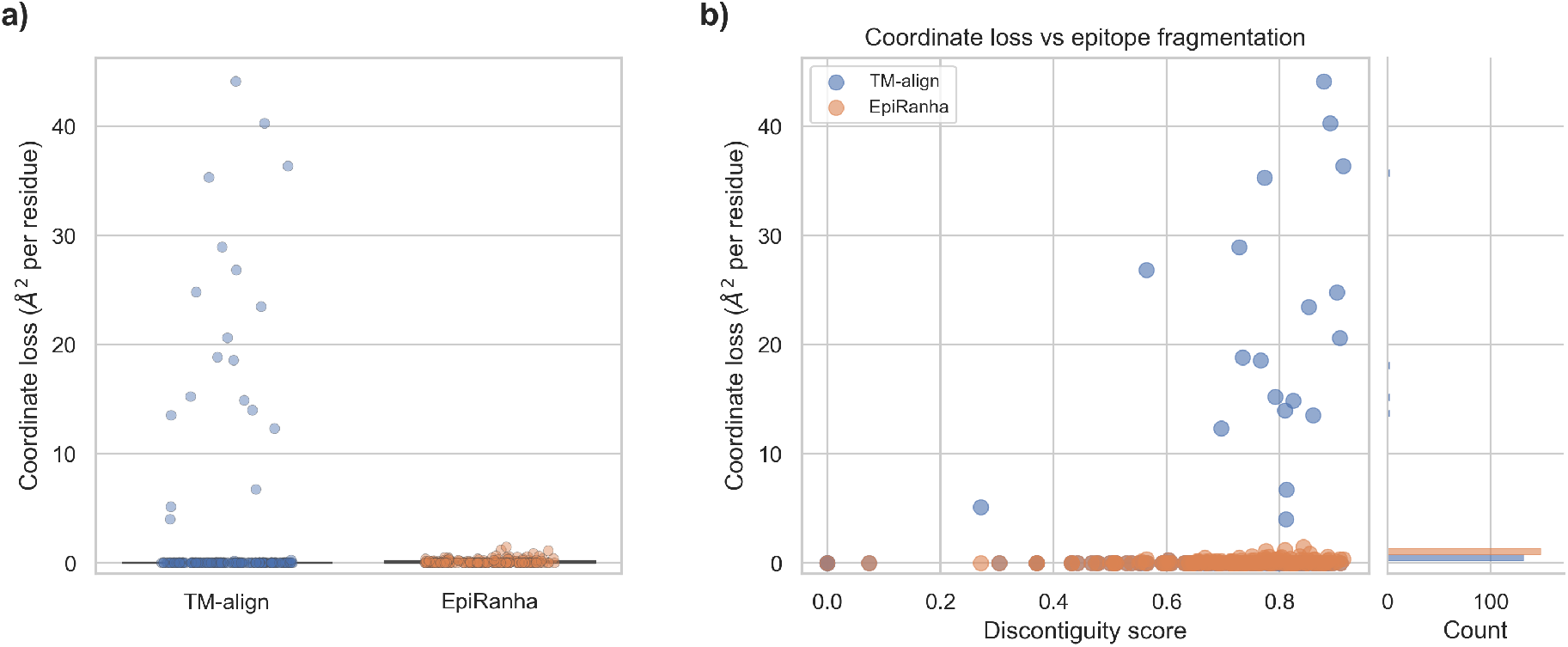
Impact of epitope discontiguity on cognate epitope–antigen alignment accuracy. (a) Distribution of coordinate loss for self-matched epitopes comparing TM-align and EpiRanha. Points indicate individual epitopes; boxes summarize the median and interquartile range. (b) Relationship between coordinate loss and epitope discontiguity. Marginal histograms show the distribution of coordinate loss for each method.

Taken together, these results demonstrate that EpiRanha not only achieves more consistent recovery of native epitopes in terms of coordinate-based loss than a rigid, structure-only baseline, but also performs well under a complementary sequence-derived (BLOSUM) loss. This indicates that EpiRanha extends beyond pure structural alignment by incorporating an additional, informative dimension of residue-level features, while maintaining strong geometric fidelity.

#### 3.1.2 Robustness to Epitope Discontiguity

To better understand the source of TM-align’s self-matching failures, we analyzed the coordinate loss as a function of epitope discontiguity. We quantified epitope discontiguity using the score defined in Section 2.4.4, which increases with both the number of gaps and the separation between discontinuous residue segments.

As shown in Figure 5b, EpiRanha maintains near-zero loss across the full range of fragmentation levels. TM-align, on the other hand, shows a clear increase in loss as discontiguity increases, indicating systematic difficulty in aligning fragmented epitopes. A more detailed characterization of TM-align failure modes, including the influence of epitope and antigen length and cross-antigen behavior, is provided in Appendix J.

This behavior likely explains the diagonal gaps in the heatmaps, indicating poor alignments (Figures 3 and 5a) where TM-align frequently recovers only a subset of epitope residues, resulting in false-negative self-matches. EpiRanha avoids this failure mode by construction, as it performs flexible, residue-level matching and always produces full-length epitope alignments. These results demonstrate that robustness to epitope discontiguity is a key factor underlying EpiRanha’s improved self-matching performance.

#### 3.1.3 Cross-Antigen Epitope Matching

Off-diagonal entries in the epitope-antigen similarity heatmaps provide insight into how each method behaves when aligning epitopes to non-cognate antigens. These matches do not correspond to known binding events, but instead reflect structural similarity of certain antigen regions to the input epitope. EpiRanha consistently identifies a larger number of low-loss matches between epitopes and non-cognate antigens than TM-align, indicating increased sensitivity to epitope-scale similarity beyond the native antigen context. TM-align, in contrast, produces comparatively sparse off-diagonal matches. Representative examples and corresponding scores from both methods are shown in Figure 6. Further interpretation of these examples, along with the overall distribution of cross-antigen losses for non-matching epitope–antigen pairs, is provided in Appendix K. Together, these examples illustrate the difference in matching behavior induced by each method’s design.

**Figure 6:**
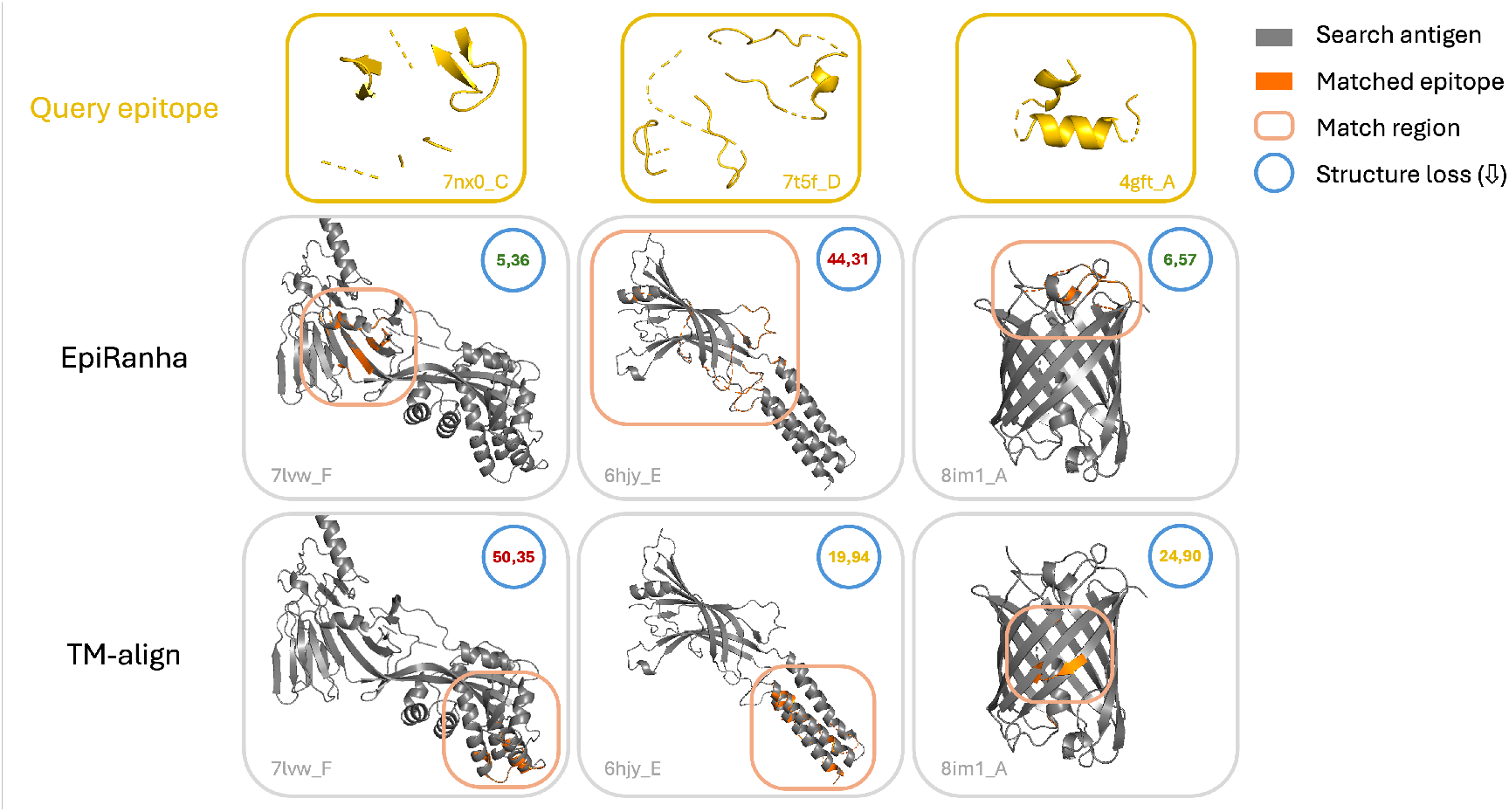
Structural visualizations of off-diagonal matching results. Three representative examples of cross-antigen matching results for a query epitope (first row) using EpiRanha (second row) and TM-align (third row). Structure loss scores (lower is better) indicate the structural quality of the match.

#### 3.1.4 Match Confidence Score

Although structure and feature losses are already a decent indication of the quality of a specific match, we also calculate a per-match confidence score for EpiRanha that represents the statistical likelihood of finding a random match with the same or better loss. Figure 7 shows these scores for three different cases. The first is cognate, i.e. self-matching, where the confidence is quite high as expected. The other two cases are cross-antigen epitope matching with low and high loss, respectively. The confidence score for cross matches with low loss is significantly higher than for those with a high loss. Some of the low-loss matches even approach the confidence of cognate matches, although the average is naturally lower. Essentially, the confidence score provides an interpretable metric for the reliability of a given match found by EpiRanha.

**Figure 7:**
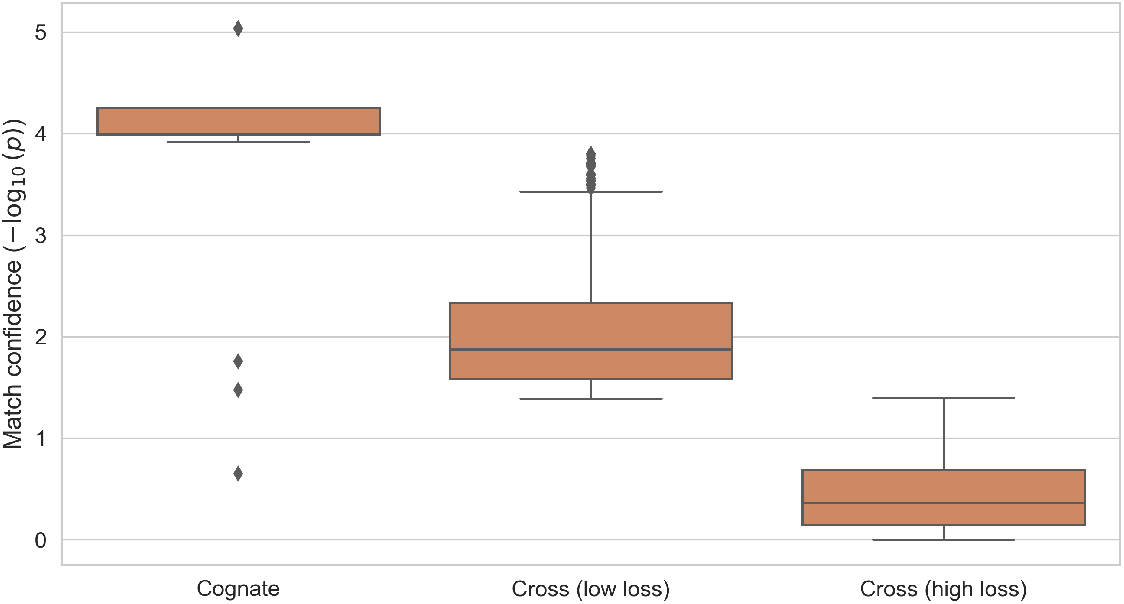
EpiRanha match confidence scores. Confidence scores, i.e. negative log of the p-value relative to a random decoy set, for three scenarios: cognate (self-matching), cross-matching with low loss ≤ 25 Å^2^, and cross-matching with high loss *>* 25 Å^2^.

### 3.2 Accuracy-Efficiency Tradeoff

While EpiRanha consistently outperforms TM-align in matching accuracy, this improvement comes at a computational cost. Figure 8 illustrates the tradeoff between coordinate loss improvement and runtime for self-matching cases. Points are colored by the reduction in coordinate loss achieved by EpiRanha relative to TM-align.

**Figure 8:**
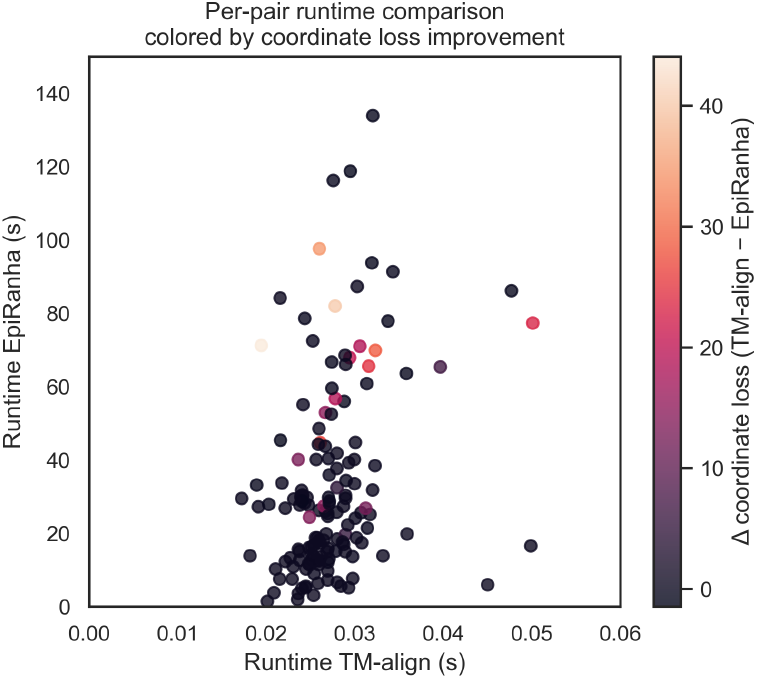
Runtime comparison between TM-align and EpiRanha, colored by structural loss improvement. Each point represents a self-matching epitope-antigen pair, with TM-align runtime on the x-axis and EpiRanha runtime on the y-axis. Points are colored by the improvement in coordinate-based loss achieved by EpiRanha, Δloss = loss_TM-align_ − loss_EpiRanha_, where positive values indicate superior performance of EpiRanha. Larger accuracy gains are associated with longer runtimes, highlighting the accuracy-efficiency tradeoff.

TM-align is substantially faster, with typical runtimes on the order of 10^−2^ seconds, compared to tens of seconds for EpiRanha under the settings used in this study. However, the largest runtime increases for EpiRanha coincide with the largest accuracy gains, likely for larger and more fragmented epitopes where TM-align struggles most. Note that TM-align was executed on a single vCPU, whereas EpiRanha was run on a multi-core instance (192 vCPUs) to enable parallel beam search. This parallelism scales with the size of the target antigen and is therefore underutilized for smaller proteins. In practice, several straightforward optimizations, such as surface-size–adaptive parallelism, could substantially reduce EpiRanha’s wall-clock runtime.

Importantly, EpiRanha’s runtime is tunable through parameters such as beam width and neighborhood size, which were fixed here to prioritize matching accuracy (see Appendix E). These results therefore highlight a clear accuracy-efficiency tradeoff: EpiRanha incurs higher computational cost, but delivers substantial and systematic gains in matching reliability, especially in challenging epitope configurations.

### 3.3 Multiple Matches in EpiRanha’s Beam Search

Beyond identifying a single best match, we show that EpiRanha can recover multiple distinct epitope-like regions on the same antigen surface for a given query epitope. These alternative matches correspond to spatially separated surface patches that exhibit comparable geometric and sequence-level consistency with the query epitope. To quantify spatial separation between these regions, candidate epitope paths produced by the beam search were grouped using residue-set clustering based on Jaccard distance between predicted residue sets (see Appendix G.1). Representative examples, quantitative separation statistics, and further details are reported in Appendix L.

## 4 Discussion

Epitope matching is a fundamental yet challenging task in computational immunology. Accurate epitope similarity scoring is critical for identifying epitope-like regions across protein surfaces, enabling the assessment of potential off-target risks and the construction of diverse training sets for downstream predictive tasks. Improving our understanding of these relationships is therefore essential for the development of more reliable computational models for epitope prediction and cross-reactivity analysis.

Here, we introduced **EpiRanha**, a hybrid sequence–structure framework that combines residue-level ESM-2 embeddings with an E(n)-equivariant graph network and a flexible beam-search matching algorithm. By integrating learned residue-level features with spatial information, EpiRanha performs residue-level matching without requiring contiguous protein segments or a single global superposition, enabling recovery of conformational epitopes that challenge traditional methods [33, 34]. Across our benchmarks, EpiRanha consistently recovers cognate epitope–antigen pairs that are missed by TM-align, which produces false-negative alignments most frequently for highly discontiguous epitopes. This failure mode likely reflects the limited structural context available for alignment: because TM-align relies on neighboring residues for secondary structure assignment and rigid-body superposition, the absence of continuous context can hinder its ability to generate accurate structural matches [18]. TM-align remains well suited for rapid, high-throughput coarse structural alignment, whereas EpiRanha is designed to operate at the residue level in settings where structural context is sparse or fragmented and additional sequence-level constraints are required to support meaningful predictions.

EpiRanha’s design also enables detection of multiple epitope-like matches on a single antigen. This capability is biologically relevant, as the presence of multiple comparable epitope-like regions on the same antigen increases the number of independent opportunities for binding over time and therefore the potential for off-target engagement [35]. Such matches may correspond either to spatially distinct surface regions or to alternative structural realizations of the same epitope, as antibody epitopes are known to exhibit conformational variability and tolerance to local structural rearrangements [4, 36]. Although a monovalent binder can occupy only one site at a time, the presence of multiple epitope-like regions provides multiple independent opportunities for binding, increasing the likelihood that the antigen is occupied by the binder over time [35, 36].

Alternative surface-based matching methods such as SurfaceID [37] were not benchmarked in this study. These approaches are designed to compare continuous surface patches using learned geometric and chemical descriptors, rather than to score similarity between arbitrary sets of epitope residues. In contrast to EpiRanha, they do not integrate residue-level sequence embeddings or perform flexible residue correspondence, which limits their sensitivity to discontiguous conformational epitopes. By jointly modeling learned sequence-level similarity and local 3D geometry within a flexible pathfinding framework, EpiRanha enables epitope similarity scoring beyond the scope of rigid structural alignment or surface fingerprint–based methods.

Runtime and computational trade-offs are an important consideration. EpiRanha is slower than TM-align due to beam-search expansion and residue-level scoring, though parallelization across multiple CPU cores and GPU acceleration can mitigate this cost. Runtime can also be tuned by adjusting beam width or neighborhood parameters, typically at the expense of some alignment quality. Further speedups are possible by constraining the search space. Although EpiRanha allows matches anywhere on the antigen structure, epitope recovery is typically only meaningful on solvent-exposed residues. Restricting beam-search expansion to a pre-defined set of surface-accessible residues (e.g., based on solvent-accessible surface area) would both reduce runtime and enforce biologically relevant matches, while preserving the generality of the current formulation.

Another useful adaptation to EpiRanha would be to allow for partial matches. By default, the algorithm is configured to return full-length epitope matches, but the search framework can be adapted to identify partial or gapped motifs by adjusting alignment thresholds and termination criteria. This flexibility enables analysis of truncated or structurally variable epitopes while preserving conservative defaults for interpretability.

For future practical deployment, we envision EpiRanha as part of a two-step epitope similarity pipeline. High-throughput global structural alignment tools such as TM-align or Foldseek can first be used to capture clear cases of overall fold similarity. EpiRanha can then be applied as a refinement step to resolve fragmented, discontinuous, or ambiguous epitope matches that are poorly handled by rigid alignment alone. Non-cognate epitope-antigen matches remain exploratory but highlight EpiRanha’s capacity to detect plausible epitope-like surfaces beyond curated immunological datasets. Future work will focus on integrating experimental cross-reactivity data, developing adaptive beam-search heuristics to further reduce runtime, and extending the framework to confidently identify partial or loop-dominated epitope motifs.

In conclusion, EpiRanha complements existing structural alignment tools by combining sequence-level context, geometric awareness, and flexible residue-level matching within a single framework. This approach allows for sensitive and interpretable discovery of both known and previously uncharacterized epitope-like regions across proteins. By doing so, it overcomes a key limitation in accurately scoring similarity between conformational epitopes, which is essential for understanding antibody cross-reactivity, while remaining practical for applications in antibody design and computational immunology.

## 5 Availability

All data used in this paper is publicly available. Code (pip-installable package), documentation, and precomputed embeddings will be published upon acceptance.

## A Related Work

Epitope similarity poses a distinct challenge for classical structural alignment because epitopes are small, frequently discontinuous surface patches embedded in larger proteins. Methods that treat each structure as a rigid body and return a single best global superposition can therefore miss partial matches and show reduced performance when aligning short segments to full-length proteins.

Traditional rigid-body methods such as TM-align quantify similarity via a length-normalised TM-score, with related approaches using iterative superposition or fragment optimisation [15]. Faster database-scale tools such as Foldseek trade some alignment accuracy for speed by approximating 3D comparisons as efficient 1D searches [17, 38]. GTalign similarly targets giga-scale exploration, but its authors report reduced performance for small peptides (<30 residues), limiting its utility for epitope matching tasks [16].

These limitations motivate representations that compare local surface regions, tolerate partial, discontinuous matches and enrich geometric information with physicochemical features. As modern datasets shift toward millions of experimentally solved or computationally predicted structures, the field has increasingly turned to machine learning models that operate beyond strict rigid-body assumptions. These approaches aim to capture functional similarity that may not be detectable through backbone geometry alone. MaSIF (molecular surface interaction fingerprinting) and its differentiable extension dMaSIF learn local geometric and physicochemical patterns from molecular surfaces, enabling prediction of ligand binding and protein-protein interfaces [22, 23, 39]. Building on the MaSIF framework, Surface ID adopts a similar surface representation and geometric deep learning architecture, but is tailored specifically for measuring protein surface similarity [37]. It represents protein surfaces as collections of local geodesic patches and learns compact numerical descriptors that capture their geometric and chemical properties, enabling efficient comparison and matching of surface regions across proteins. Similarly, Folddisco indexes position-independent structural motifs, such as side-chain orientations or torsion patterns, and applies rarity-based scoring to identify functionally informative motifs across large structural databases [20]. Collectively, these approaches highlight how reducing protein structures to surfaces or motifs can uncover functional relationships that remain hidden to sequence- or backbone-based comparisons. However, despite capturing global geometry and recurring physicochemical patterns, surface- and motif-based representations can miss residue-level binding details critical for molecular recognition. To address this, graph-based approaches have recently emerged as a powerful framework for representing protein structures at residue-level resolution, enabling fine-grained similarity assessment as well as efficient comparison, clustering, and design.

Early frameworks such as GraphPPIS [40] and DeepRank-GNN [41] used GNN architectures to model protein-protein interfaces, demonstrating that accurately capturing spatially neighboring residues is essential for interaction prediction. In the context of protein design, ProteinSolver [42] applied graph convolutional networks to reconstruct sequences compatible with predefined folds using constraint satisfaction, while subsequent studies further highlighted the versatility of GNNs for integrating sequence, structure, and function information [43] and for tasks such as binding-affinity prediction [44]. Within this broader class of methods, equivariant graph neural networks have received particular attention. Satorras et al. [30] introduced the E(n)-Equivariant Graph Neural Network (EGNN), which maintains equivariance to translations, rotations, reflections, and permutations, ensuring that predictions remain consistent under these geometric transformations. In contrast to earlier architectures, the EGNN achieves this without relying on computationally expensive higher-order tensor representations, while still delivering competitive or superior performance on tasks such as molecular property prediction. Building on this framework, recent work from Greener et Al. [45] has extended equivariant GNNs to protein comparison tasks by training models that embed protein domains into low-dimensional vectors using supervised contrastive learning guided by curated structural classifications. These embeddings capture biologically meaningful relationships between folds and enable rapid comparison across large structure databases by simple cosine similarity, transforming equivariant GNNs from molecular modeling tools into scalable solutions for protein structure search. While graph-based architectures excel at representing the geometric and relational aspects of protein structures, sequence-informed models offer a complementary perspective by extracting evolutionary and biochemical signals directly from sequence data.

For instance, Lin et al. [24] introduced ESM-2, an LLM trained on millions of protein sequences, able to generate residue-level embeddings that encode both evolutionary and physicochemical information. Building on this progress, recent work has increasingly explored combining sequence-derived embeddings from LLMs with graph-based representations of protein structure. Integrating multi-modal data allows models to leverage complementary sources of information, capturing both local structural context and global sequence-level features. Blaabjerg et al. [46] demonstrated this with SSEmb, which integrates structural data and multiple sequence alignments to predict protein variant effects, achieving robustness in settings with limited sequence information. Although a wide range of graph-based, sequence-based, and hybrid machine learning models exists for general protein representation and interaction prediction, several specialised tools have recently been developed for B-cell epitope prediction.

These models aim to identify antigenic determinants that mediate antibody recognition and are therefore essential for therapeutic and vaccine design. Recent examples include DiscoTope [33], which leverages inverse folding latent structure representations to improve conformational B-cell epitope prediction; BepiPred-3.0 [34], a sequence-based predictor that exploits protein language model embeddings to enhance accuracy on both linear and conformational epitopes; AbEpiTope-1.0 [47], which integrates AlphaFold-Multimer modeling with inverse folding to perform antibody-specific B-cell epitope prediction; and GraphBepi [25], which fuses predicted AlphaFold-2 structures with ESM-2 embeddings.

By contrast, approaches for epitope similarity scoring, that is, comparing epitopes across proteins to assess potential cross-reactivity or shared recognition, remain far less developed. Whereas prediction methods aim to identify candidate epitopes within a single antigen, similarity scoring seeks to quantify relationships between epitopes across different antigens, a capability critical for studying immune cross-reactivity, antibody specificity, and off-target effects. This distinction highlights a gap in current methodology and underscores the need for complementary frameworks that move beyond epitope identification toward systematic comparison.

## B Dataset Construction and Preprocessing

To ensure high-quality structural data for the nanobody-antigen complexes sourced from SAbDab-nano, we set a structural resolution cutoff of ≤ 3Å. Duplicates were removed, and only complexes consisting of a single-chain antigen and a single-chain nanobody of camelid origin (camel or llama) were retained. Epitope and paratope residues were defined based on inter-protein contacts, using a C_*α*_-C_*α*_ distance cutoff of 4Å. Complexes lacking any contacts within this threshold were excluded from further analysis.

To reduce redundancy in the dataset, hierarchical clustering was performed on epitope centroids using a cutoff of 10Å for the inter-cluster distance. From each resulting cluster, a single representative complex was selected at random. Finally, complexes in which the same nanobody was observed to bind multiple distinct antigens were discarded.

Finally, the structures were preprocessed by removing duplicates from the PDB file, removing waters and heterogens, fixing gaps and renumbering consecutively. Subsequently, the structure is relaxed through minimization of its free energy. Processing steps were done using OpenMM [48], and a summary of these steps is illustrated in Figure B1. From the processed complexes, we only retained the antigens, as we are focusing on epitope matching independently of the nanobody interaction. After this step, we removed duplicate antigens in the data to reduce model overfitting. It is important to note that these antigens taken directly out of complexes correspond to their holo (bound) states, which may introduce biases compared to apo (unbound) antigens [49].

For representation learning purposes, we augmented the SabDab-nano dataset with AlphaFold-predicted surface proteins. These were not further processed.

**Figure B1:**
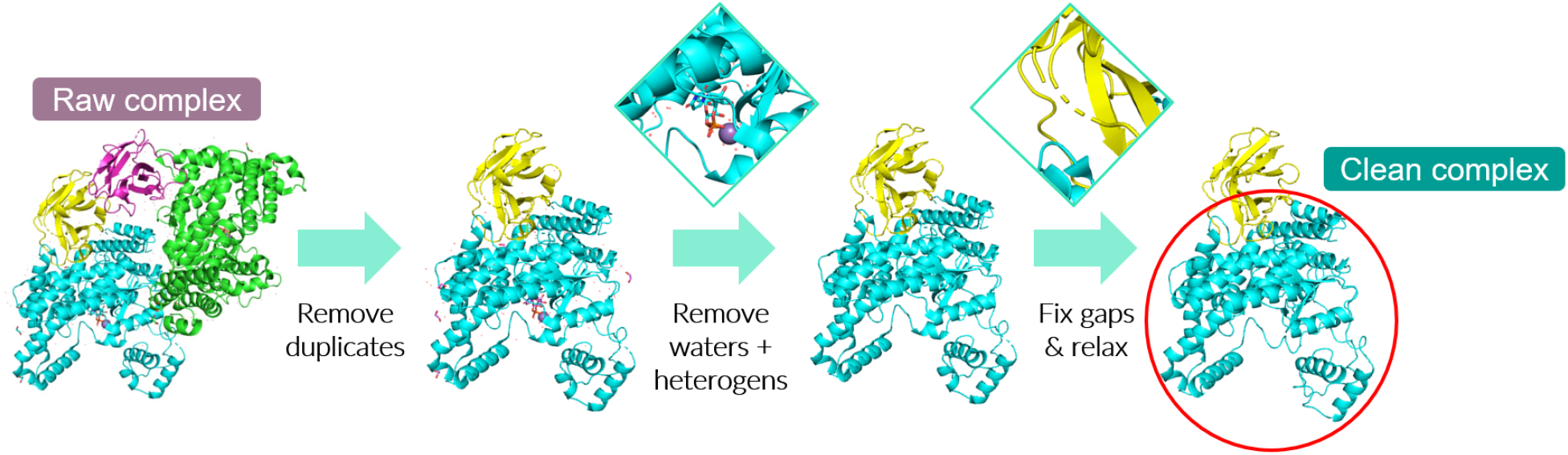
Preprocessing steps applied to each raw nanobody-antigen complex sourced from SAbDab-nano, ultimately resulting in a clean complex from which we only retain the antigen.

## C EGNN Workflow and Training

A complete overview of the EGNN workflow used in EpiRanha, from protein to the generation of per-residue embeddings, is depicted in Figure C1.

**Figure C1:**
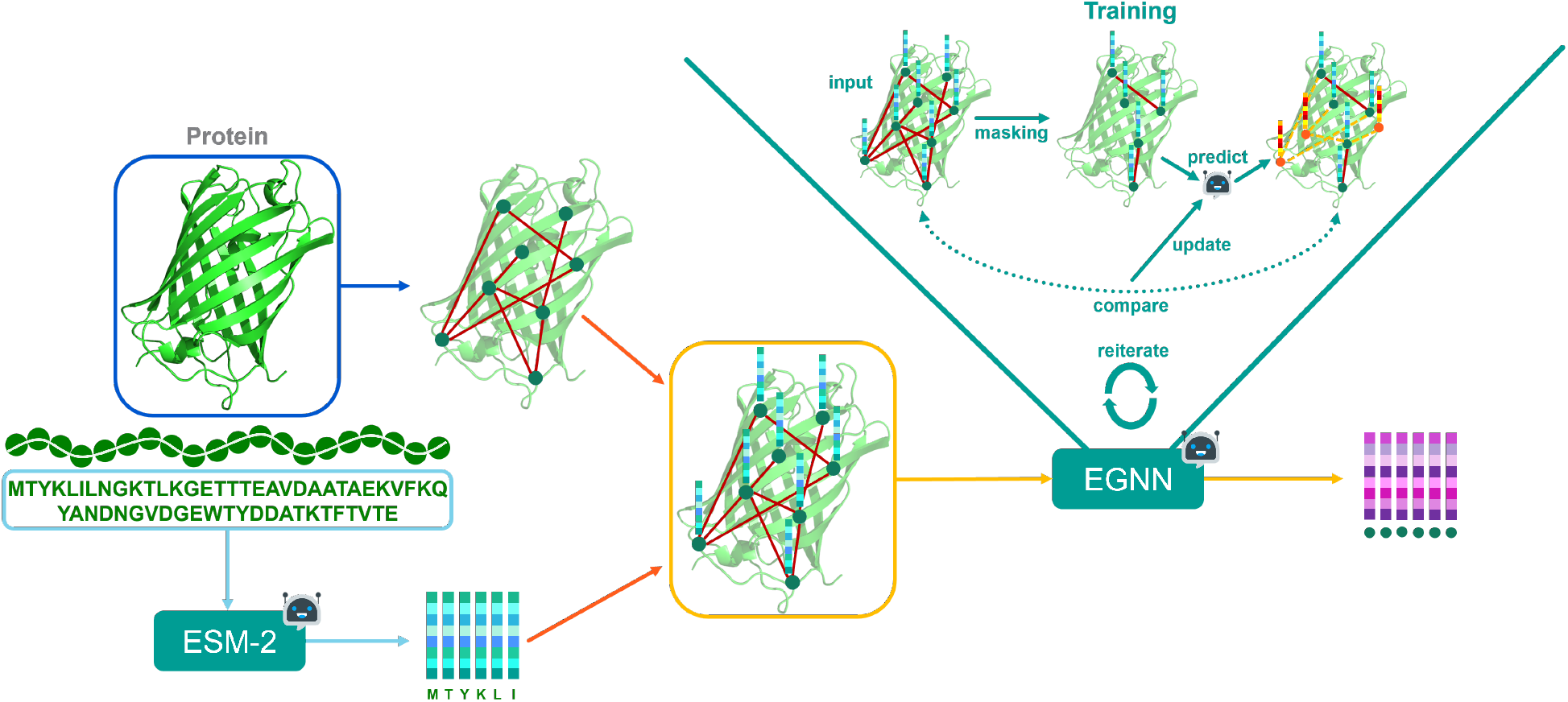
Overview of the EGNN workflow in EpiRanha. The protein sequence is first processed through ESM-2 to generate per-residue embeddings. These embeddings, along with the C_*α*_ atomic coordinates, form the input graph for the EGNN. During training, masking is applied to randomly selected residues which requires the EGNN to reconstruct them using message passing from neighboring nodes. After multiple epochs, the final per-residue embeddings are extracted for use in epitope matching tasks. Robot icons represent ML-based components.

### C.1 Training Procedure

EGNN training follows a three-stage process: hyperparameter optimization, model validation, and final embedding generation. First, hyperparameter optimization is performed to identify suitable architectural and optimization settings. Bayesian optimization with Optuna [50] is used to explore the hyperparameter space. The data were split into 70% for training, 20% for validation, and 10% for testing. Candidate configurations are evaluated based on validation loss after a fixed number of training epochs. The evaluated hyperparameters, their respective search ranges, and the optimal values are summarized in Table C1. Unless stated otherwise, all remaining parameters were held fixed across experiments.

Second, model stability and generalization are assessed through cross-validation using the selected hyperparameter configuration. This step is used to verify robustness but does not directly affect the final embedding generation. Finally, a single model per configuration is trained to convergence and used to generate embeddings for all proteins included in the study.

All models were trained for a maximum of 500 epochs, with model selection based on the lowest validation loss and no early stopping. A fixed random seed (42) was used to ensure reproducibility. Optimization was performed using the Adam optimizer with weight decay, and a step-based learning-rate schedule was applied.

### C.2 Loss Function

The EGNN is trained using a composite reconstruction loss consisting of a coordinate reconstruction term and a feature reconstruction term.

The coordinate loss measures the mean squared error between predicted and true C_*α*_ coordinates, encouraging accurate geometric reconstruction. The feature loss measures the mean squared error between predicted and original ESM-2 embeddings, encouraging preservation of sequence-derived information.

The total loss of the model is computed as a weighted sum of the coordinate and feature losses:

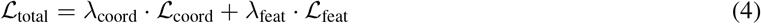

To balance the scale of the two losses, we set the weights to *λ*_coord_ = 0.04 and *λ*_feat_ = 0.96, which ensures that both contribute comparably to the total loss. These values were chosen empirically but could be tuned further for optimization.

**Table C1:** Summary of tested hyperparameters, their respective search ranges, and the optimal values obtained from the EGNN hyperparameter search. Continuous values were sampled for parameters with specified floating-point ranges, while integer values were sampled for the others. The learning rate was adjusted using a step-based schedule, reducing it every ‘step size’ epochs by ((1 − *γ*) *×* 100)%.

| Hyperparameter | Tested Range | Optimal Value |
| --- | --- | --- |
| learning rate | $[10^{-5}, 10^{-2}]$ | $1.04 \times 10^{-3}$ |
| weight decay | $[10^{-6}, 10^{-3}]$ | $6.95 \times 10^{-6}$ |
| step size | [5, 20] | 12 |
| gamma ($\gamma$) | [0.8, 0.95] | 0.93 |
| batch size | [1, 6] | 4 |
| model depth | [4, 12] | 9 |
| hidden dimension size | [8, 32] | 16 |
| nr of nearest neighbors | [2, 32] | 5 |
| nr of fourier features | [2, 5] | 4 |
| coord weights clamp value | [0.5, 5.0] | 1.49 |
| dropout | [0.0, 0.5] | 0.14 |

### C.3 Masking Strategy and Dataset Composition

Masking was applied at the residue level by removing both coordinates and ESM-2 embeddings for selected residues during training. The masking fraction, which determines the proportion of residues hidden during training, was not optimized through the hyperparameter search. Because lower masking inherently reduces reconstruction loss, it was instead evaluated empirically based on downstream performance. Four masking fractions (0.05, 0.1, 0.15, and 0.3) were preselected, resulting in separate trained models for each setting.

Two datasets were used to train the models: one containing only selected antigens from SAbDab-nano and another that combined SAbDab-nano with AlphaFold 2 surface proteins. This led to eight different model variants, each trained with a different combination of masking fraction and dataset composition, that produced eight unique embeddings for each protein.

### C.4 Cross-Validation and Embedding Generation

For cross-validation, 90% of the data were used for training and 10% for testing, with alternating validation splits across folds. Final models used for embedding generation were trained on 90% of the data, reserving 10% for validation to maximize information from the full dataset. Since the EGNN does not directly perform epitope matching but rather encodes information into embeddings, concerns about data leakage were not an issue.

Each of the eight EGNN variants was assessed using five-fold cross-validation on a common held-out test set. For the final embedding generation, models were retrained without cross-validation to ensure that all proteins from SAbDab-nano and AlphaFold 2 contributed to the embedding space.

## D EGNN Reconstruction Performance

Although the EGNN is not evaluated as a standalone task in this work, we assess its reconstruction performance to verify stable training and meaningful representation learning prior to downstream epitope matching.

### D.1 Quantitative Reconstruction Performance

Reconstruction performance was evaluated using the validation loss and independent reconstruction metrics across five cross-validation folds. We report the total reconstruction loss, the coordinate and feature reconstruction loss, and the cosine similarity between reconstructed and original ESM-2 embeddings. Results are summarized in Tables D1 and D2.

Across all masking fractions, models trained on the combined SAbDab–AlphaFold dataset consistently achieved lower validation loss and higher embedding similarity than models trained on SAbDab alone, indicating improved stability and generalization when incorporating structurally diverse AlphaFold data. Increasing the masking fraction led to higher reconstruction error, reflecting the increased difficulty of the task, while preserving consistent relative performance trends across datasets.

**Table D1:** Cross-validation results per fold for different masking fractions using an EGNN trained on solely antigens. The best average values across masking fractions are bolded.

| Masking Fraction | Fold | Loss (↓) | | | Feature Cosine Similarity (↑) |
| --- | --- | --- | --- | --- | --- |
|  |  | Total | Coordinate | Feature |  |
| 0.05 | 1 | 0.1055 | 1.6995 | 0.0391 | 0.6827 |
|  | 2 | 0.1083 | 1.8275 | 0.0366 | 0.6966 |
|  | 3 | 0.1328 | 2.4161 | 0.0377 | 0.6883 |
|  | 4 | 0.1186 | 2.0697 | 0.0373 | 0.6936 |
|  | 5 | 0.0966 | 1.5211 | 0.0372 | 0.6911 |
|  | **Avg** | **0.1124** | **1.9068** | **0.0376** | **0.6905** |
| 0.10 | 1 | 0.1635 | 3.1771 | 0.0380 | 0.6842 |
|  | 2 | 0.1537 | 2.9429 | 0.0375 | 0.6874 |
|  | 3 | 0.0998 | 1.5881 | 0.0378 | 0.6851 |
|  | 4 | 0.1581 | 3.0527 | 0.0375 | 0.6877 |
|  | 5 | 0.0943 | 1.4592 | 0.0375 | 0.6886 |
|  | **Avg** | 0.1339 | 2.4440 | 0.0377 | 0.6866 |
| 0.15 | 1 | 0.1725 | 3.4002 | 0.0381 | 0.6836 |
|  | 2 | 0.1357 | 2.4956 | 0.0382 | 0.6810 |
|  | 3 | 0.1273 | 2.2722 | 0.0379 | 0.6822 |
|  | 4 | 0.1655 | 3.2262 | 0.0380 | 0.6828 |
|  | 5 | 0.1570 | 3.0207 | 0.0377 | 0.6858 |
|  | **Avg** | 0.1516 | 2.8790 | 0.0380 | 0.6831 |
| 0.30 | 1 | 0.1615 | 3.1254 | 0.0380 | 0.6839 |
|  | 2 | 0.1846 | 3.6947 | 0.0383 | 0.6808 |
|  | 3 | 65.6894 | 1640.2267 | 0.0837 | 0.6523 |
|  | 4 | 0.1719 | 3.3954 | 0.0376 | 0.6863 |
|  | 5 | 0.2390 | 5.0697 | 0.0377 | 0.6856 |
|  | **Avg** | 13.2893 | 331.1024 | 0.0471 | 0.6778 |

Importantly, absolute reconstruction loss values are not interpreted as performance targets, but rather as indicators of successful and stable training under varying masking regimes.

### D.2 Qualitative Reconstruction Behavior

To complement the quantitative results, Figure D1 provides qualitative examples of reconstructed residue coordinates under different masking fractions. At low masking levels, predicted coordinates closely follow the native backbone geometry. As the masking fraction increases, reconstructions become progressively less constrained and may exhibit localized distortions, reflecting the reduced contextual information available during message passing.

These qualitative trends are consistent with the observed increase in reconstruction loss and illustrate the trade-off between reconstruction difficulty and representation expressiveness.

## E Design Choices and Robustness Analyses

This appendix provides extended qualitative analyses that informed the design choices of EpiRanha, including embedding training data composition, masking fractions, and beam search parameter selection. All analyses were conducted on representative query-search protein combinations and aim to assess robustness and behavioral consistency rather than optimize downstream performance.

**Table D2:** Cross-validation results per fold for different masking fractions using an EGNN trained on both antigens and AlphaFold data. The best average values across masking fractions are bolded.

| Masking Fraction | Fold | Loss (↓) | | | Feature Cosine Similarity (↑) |
| --- | --- | --- | --- | --- | --- |
|  |  | Total | Coordinate | Feature |  |
| 0.05 | 1 | 0.1049 | 1.7892 | 0.0348 | 0.7104 |
|  | 2 | 0.1078 | 1.8558 | 0.0350 | 0.7085 |
|  | 3 | 0.0766 | 1.0901 | 0.0344 | 0.7141 |
|  | 4 | 0.1164 | 2.0591 | 0.0355 | 0.7033 |
|  | 5 | 0.1112 | 1.9338 | 0.0353 | 0.7054 |
|  | **Avg** | **0.1034** | **1.7456** | **0.0350** | **0.7083** |
| 0.10 | 1 | 0.1394 | 2.6040 | 0.0367 | 0.6907 |
|  | 2 | 0.1129 | 1.9724 | 0.0354 | 0.7036 |
|  | 3 | 0.1118 | 1.9518 | 0.0351 | 0.7070 |
|  | 4 | 0.0824 | 1.2103 | 0.0354 | 0.7044 |
|  | 5 | 0.0916 | 1.4500 | 0.0350 | 0.7087 |
|  | **Avg** | 0.1076 | 1.8377 | 0.0355 | 0.7029 |
| 0.15 | 1 | 0.1196 | 2.1364 | 0.0356 | 0.7025 |
|  | 2 | 0.1148 | 2.0250 | 0.0352 | 0.7058 |
|  | 3 | 0.1210 | 2.1686 | 0.0357 | 0.7010 |
|  | 4 | 0.1184 | 2.1144 | 0.0352 | 0.7059 |
|  | 5 | 0.1164 | 2.0678 | 0.0351 | 0.7071 |
|  | **Avg** | 0.1181 | 2.1024 | 0.0354 | 0.7044 |
| 0.30 | 1 | 0.1391 | 2.6035 | 0.0364 | 0.6942 |
|  | 2 | 0.1377 | 2.5764 | 0.0360 | 0.6978 |
|  | 3 | 0.1369 | 2.5562 | 0.0361 | 0.6972 |
|  | 4 | 0.1316 | 2.4202 | 0.0363 | 0.6952 |
|  | 5 | 0.1324 | 2.4411 | 0.0362 | 0.6963 |
|  | **Avg** | 0.1355 | 2.5195 | 0.0362 | 0.6961 |

### E.1 Beam Search Parameter Selection

Beam search parameters were selected based on a qualitative trade-off between alignment quality and computational efficiency. Masking fractions, numbers of spatial neighbors, and beam widths were explored on a fixed set of representative query–search protein combinations spanning small, medium, and large epitopes.

Figure E1 summarizes these experiments. Across all tested cases, a masking fraction of 0.1 consistently produced more stable and informative embeddings than lower values, which appeared to under-regularize the model, or higher values, which degraded reconstruction quality. While masking fraction did not affect runtime, it strongly influenced alignment behavior.

Notably, embeddings trained on the combined dataset of experimental and AlphaFold-derived structures consistently exhibited lower loss scales across all tested parameter configurations compared to embeddings trained on experimental data alone. This behavior indicates improved representational stability and robustness, and it directly motivated the use of the combined training dataset for all downstream epitope matching experiments.

The number of spatial neighbors controlled the breadth of the search space. Too few neighbors resulted in brittle search behavior, particularly for larger proteins, whereas increasing this parameter improved alignment quality at the cost of additional computation. In contrast, beam width primarily affected runtime, with diminishing returns beyond moderate values.

Based on these observations, a masking fraction of 0.1, 12 neighbors, and a beam width of 500 were selected as default parameters. These settings provide a robust compromise between runtime and matching quality and are used throughout the main experiments.

**Figure D1:**
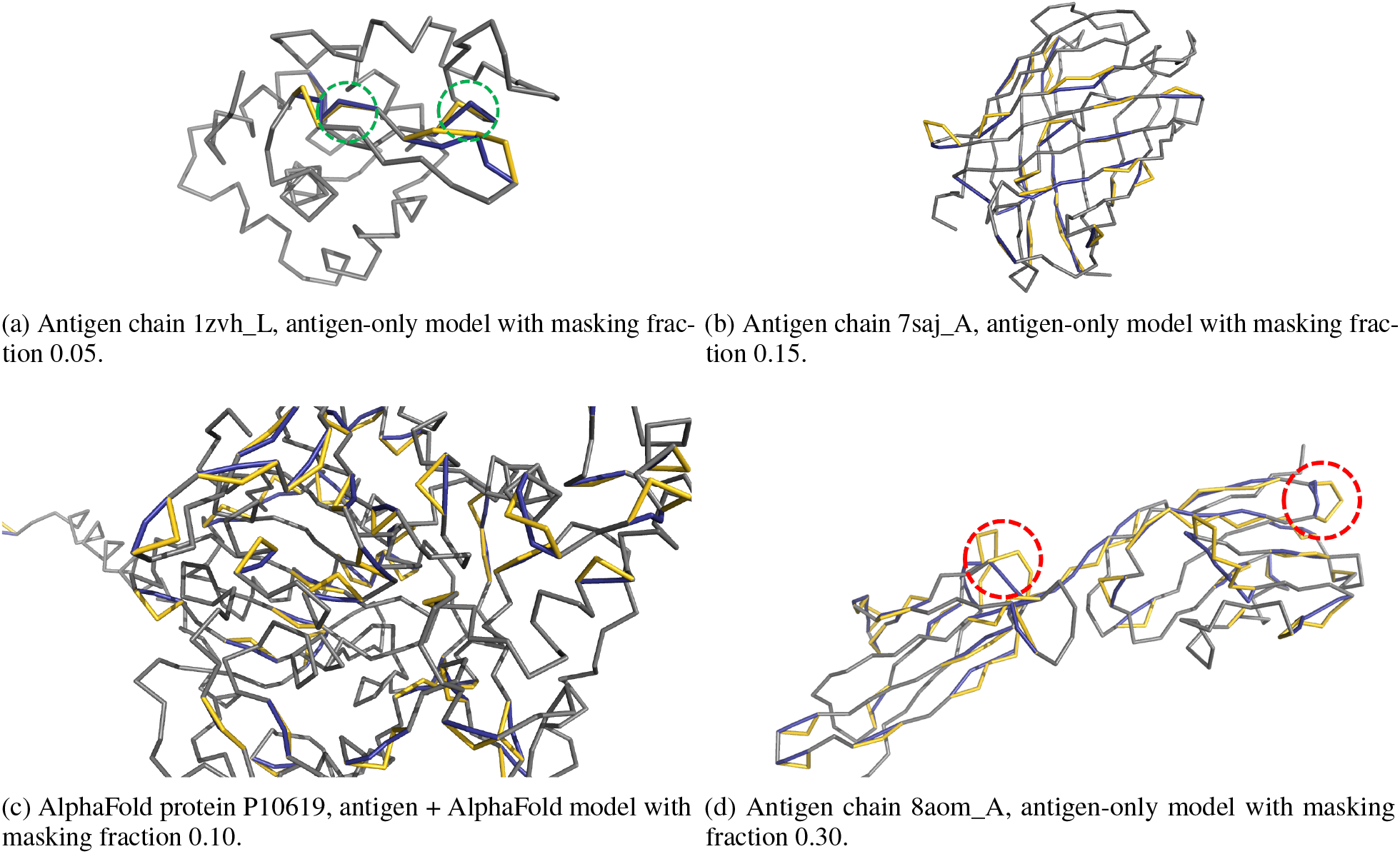
Visualization of predicted versus true residue positions across different model settings. All models shown correspond to the second fold best model, as these were the most stable across settings. Colors: Gray = structure with missing residues (model input), Gold = true masked residues, Blue = predicted coordinates. (a) Minimal masking allows the model to reconstruct missing residues with high accuracy, with some almost perfectly predicted residues highlighted in green. (b) Increased masking makes the reconstruction task more challenging. (c) A model trained on both antigens and AlphaFold proteins results in improved prediction stability. (d) Excessive masking leads to poor reconstruction, with some misaligned predictions highlighted in red.

### E.2 Modality Ablations

To assess the individual contributions of structural information and learned residue representations, we performed a series of modality ablations in which the beam search was executed using different combinations of coordinates and embeddings. These experiments serve as sanity checks to verify that EpiRanha’s performance does not arise trivially from either geometric alignment alone or sequence-derived features alone.

An overview of all evaluated model variants is provided in Table E1. In addition to the full EpiRanha configurations, we considered beam search using raw ESM-2 embeddings with and without coordinates, EGNN embeddings without coordinates, and a coordinate-only variant without any feature information. TM-align is included solely as a visual reference baseline and is not quantitatively compared in this appendix.

We consider the most straightforward validation scenario, in which an epitope is matched against the antigen from which it was derived. In this setting, a correct method should consistently recover the original epitope location. As shown in Figure E2, models that combine EGNN embeddings with coordinate-based search reliably identify the correct epitope region, producing matches with low feature and coordinate loss. In contrast, using only coordinates or only embeddings leads to less stable behavior, with increased variance and occasional misalignment. Raw ESM-2 embeddings, even when combined with coordinates, fail in some cases to fully reconstruct the original epitope, indicating that sequence-derived features alone are insufficient to capture the relevant structural context.

In contrast, models that rely exclusively on a single modality exhibit less reliable behavior. Depending on the specific query, beam search using only embeddings or only coordinates fails to recover the correct epitope location. For example, in the 8im1_A case, embedding-only variants and raw ESM-2 embeddings do not achieve the optimal match, whereas in the 3g9a_A and 6h15_B cases, multiple single-modality configurations, including coordinate-only search and embedding-only variants, fail to identify the correct epitope region. Although these examples are qualitative, this pattern was consistently observed across the evaluated dataset, suggesting that reliance on a single information source leads to fluctuating and query-dependent behavior.

**Figure E1:**
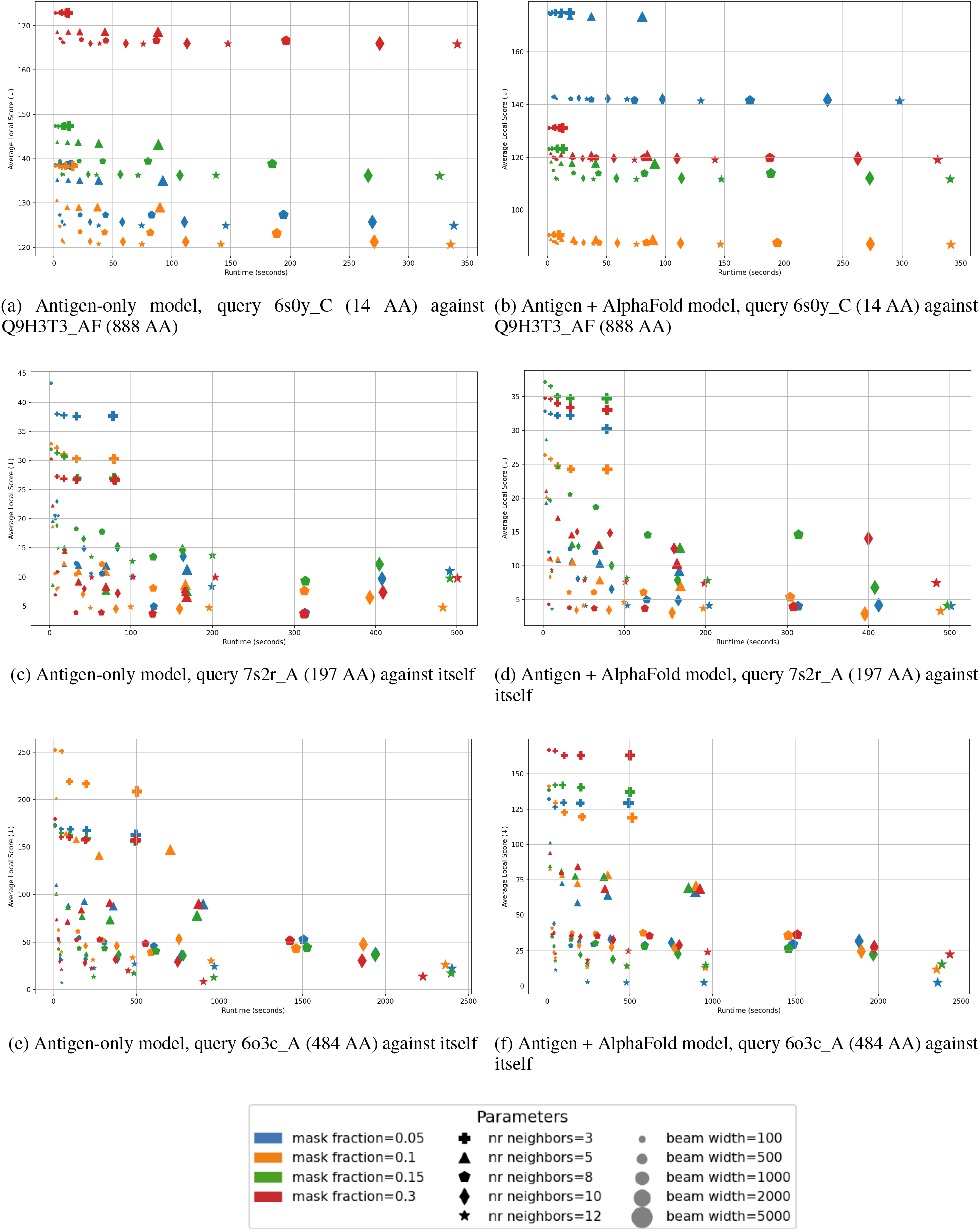
Benchmark results comparing beam search performance versus runtime for different query-search protein pairs. The left column corresponds to beam search results with embeddings from models trained on antigens only, while the right column corresponds to models trained on both antigens and AlphaFold proteins. Each row represents a different query-search protein combination, ordered by increasing epitope size. The amount of amino acids (AA) per protein is added in each caption.

**Table E1:** Description of the different model types used in the evaluation. The model type column specifies the name used in the evaluation plots. The beam search input column describes whether both coordinates and embeddings, only embeddings, or only coordinates were used during beam search. The embedding model indicates whether the model uses ESM-2-based sequence embeddings or EGNN-based structure-aware embeddings. The EGNN training data column specifies whether the model was trained on antigens only or also included AlphaFold-predicted structures.

| Model Type | Beam Search Input | Embedding Model | EGNN Training Data |
| --- | --- | --- | --- |
| ESM-2 | Coordinates + Embeddings | ESM-2 | - |
| only_feats_ESM-2 | Only Embeddings | ESM-2 | - |
| egnn_ag_only | Coordinates + Embeddings | EGNN | Antigens only |
| egnn_ag_af | Coordinates + Embeddings | EGNN | Antigens + AlphaFold |
| only_feats_egnn_ag_only | Only Embeddings | EGNN | Antigens only |
| only_feats_egnn_ag_af | Only Embeddings | EGNN | Antigens + AlphaFold |
| only_coords | Only Coordinates | - | - |
| tmalign | - | - | - |

Across all examined scenarios, models that rely exclusively on coordinates or exclusively on embeddings exhibit unstable behavior that depends strongly on the specific query–search combination. Raw ESM-2 embeddings perform well in some cases but lack the structural grounding required for reliable epitope reconstruction, even when combined with spatial search. EGNN embeddings, particularly those trained on both experimental antigens and AlphaFold-predicted structures, consistently yield more stable and accurate matches when used together with coordinate-based beam search. Taken together, these ablation studies confirm that EpiRanha’s performance arises from the integration of sequence-aware embeddings and explicit spatial information, motivating the use of this combined configuration in all downstream experiments.

## F Epitope Matching Algorithm and Pseudocode

A schematic overview of the epitope matching process in EpiRanha is shown in Figure F1.

### F.1 Path Scoring and Similarity Function

Candidate paths are evaluated using an assignment-based similarity score. Given a partial or complete path *P* in the search antigen and the query epitope *Q*, an optimal one-to-one residue correspondence is first computed by solving a linear sum assignment problem. This problem is solved using a modified Jonker-Volgenant algorithm without initialization [31], ensuring an optimal matching between residues in *P* and residues in *Q*.

Based on this residue mapping, two distance-based terms are computed. The structural term measures the sum of Euclidean distances between the three-dimensional coordinates of matched residues. The feature term measures the sum of Euclidean distances between the corresponding residue embeddings. The similarity score is defined as

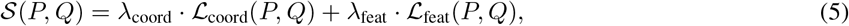

where ℒ_coord_ denotes the structural distance term, ℒ_feat_ the feature distance term, and *λ*_coord_ and *λ*_feat_ are weighting coefficients that balance both contributions. The weights were calibrated by first evaluating each term independently to assess its scale and then adjusting them such that both terms contribute comparably to the total score. Lower scores indicate higher similarity.

### F.2 Beam Search Algorithm

Algorithm 1 provides high-level pseudocode for the beam search procedure used to identify epitope-like regions in search antigens. The algorithm operates on precomputed per-residue embeddings and coordinates and incrementally constructs candidate matches by expanding spatially adjacent residues.

**Figure E2:**
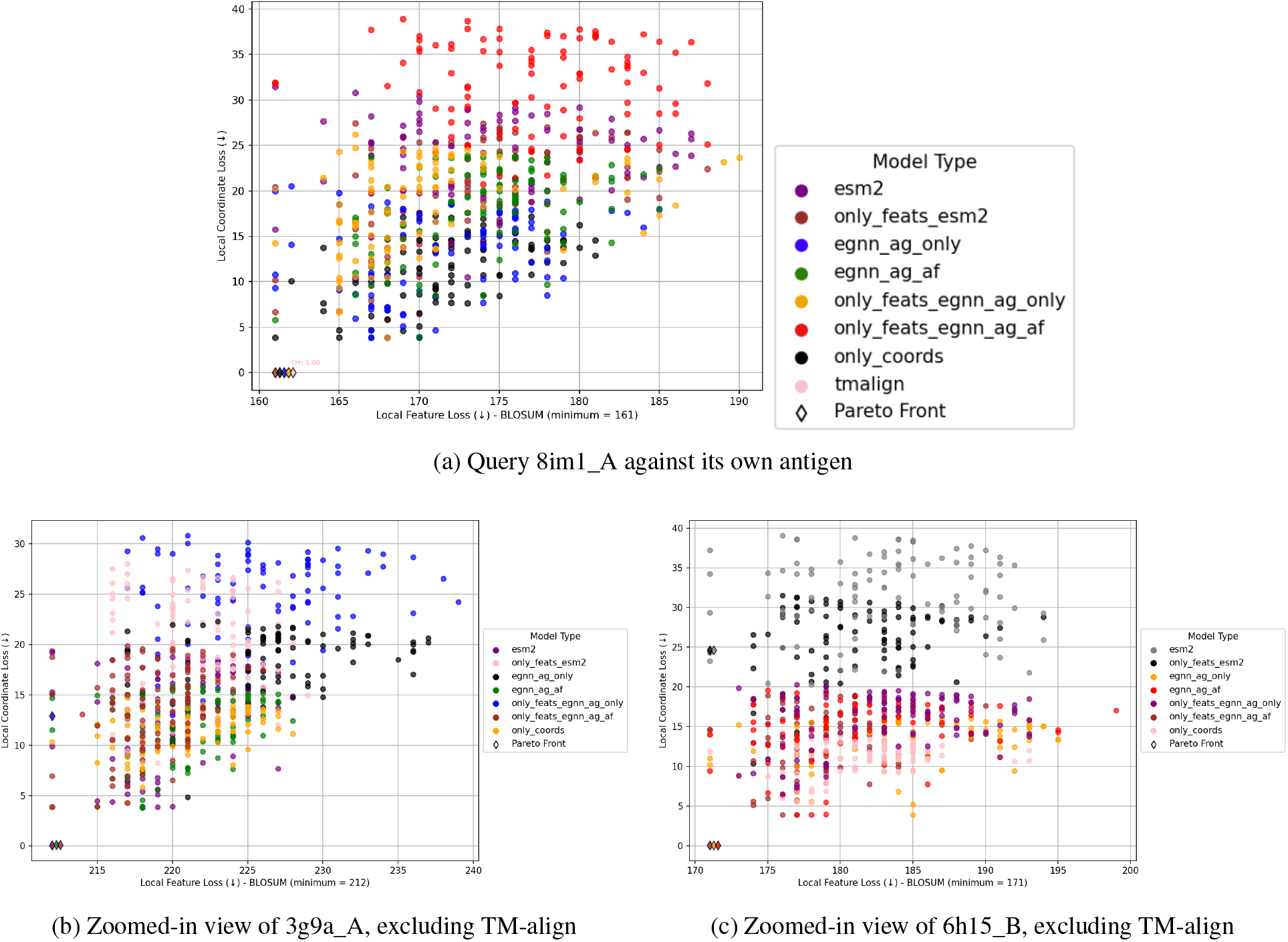
Feature loss (BLOSUM-based) versus coordinate loss for epitopes matched against their own antigens. Each plot shows the distribution of the top 100 beam search matches for different model variants. Optimal matches lie in the lower-left region of the plots. While several configurations approach the optimum, models combining EGNN embeddings with coordinate-based search consistently yield the most stable and near-optimal alignments, whereas embedding-only and coordinate-only variants frequently fail to recover the exact epitope location.

### F.3 Parameter Settings

The final parameter values used in all experiments were selected based on empirical sensitivity analyses and sanity checks described in Appendix E. Only parameters relevant to the final epitope matching procedure are listed in Table F1 for completeness.

The beam width and neighborhood size control the trade-off between computational efficiency and search completeness, while the loss weights determine the relative influence of structural versus feature similarity. Different feature representations require different weightings due to differences in scale and semantic meaning of their embedding spaces. These weights were fixed across all experiments once selected.

**Figure F1:**
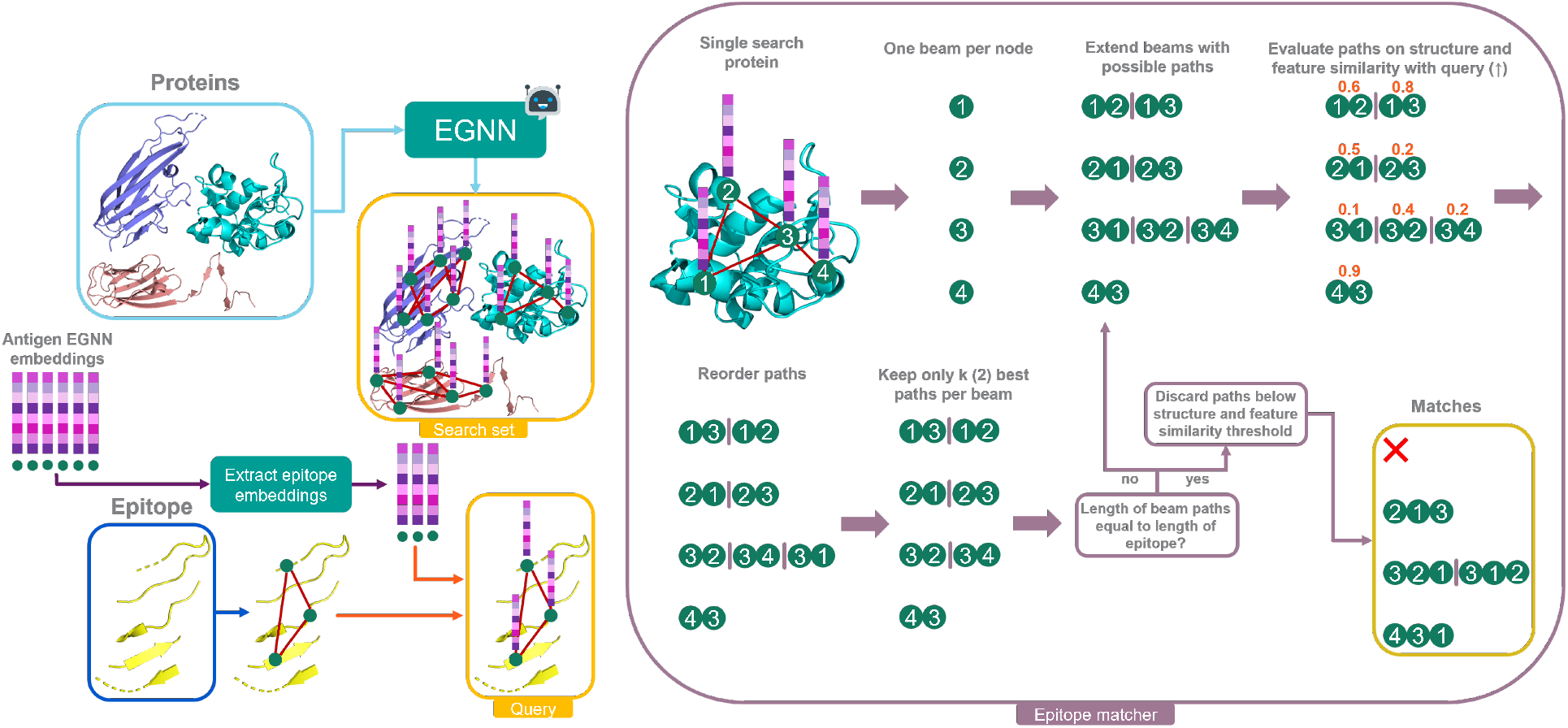
Overview of the epitope matching process using beam search in EpiRanha. The query epitope with its EGNN embeddings is extracted from an antigen and compared against a search set of proteins processed through EGNN. A beam search is initiated from each residue in the search protein, iteratively expanding paths by adding nearby residues in 3D space. Paths are ranked based on their structural and feature similarity to the query. Once paths reach the same length as the epitope, the highest-scoring matches are retained. Robot icons represent ML-based components.

#### Algorithm 1 Beam-Search–Based Epitope Matching in EpiRanha

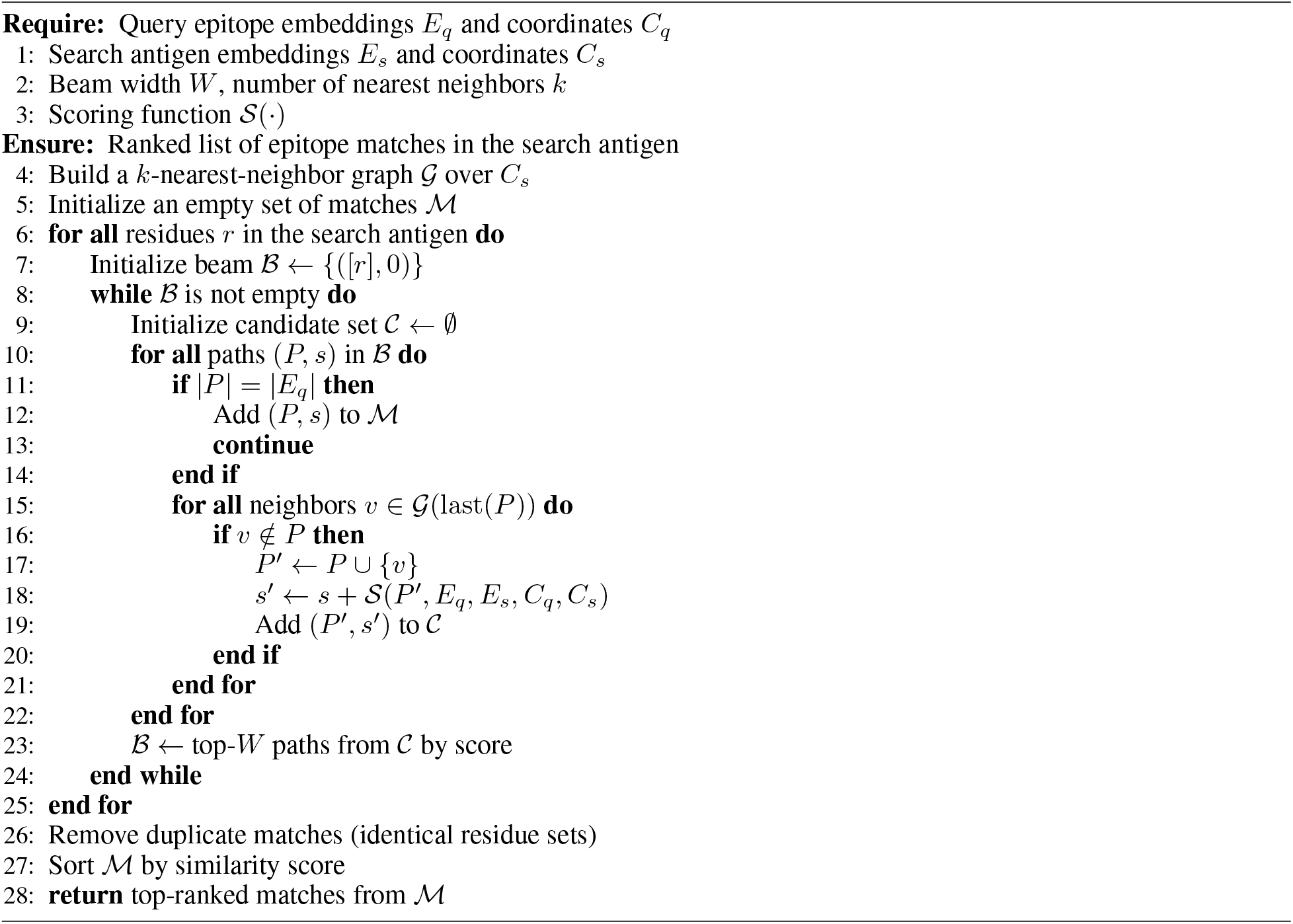

**Table F1:** Final beam search and scoring parameters used in EpiRanha.

| Parameter | Value |
| --- | --- |
| Beam width ($W$) | 500 |
| Number of nearest neighbors ($k$) | 12 |
| Structural weight ($\lambda_{\text{coord}}$, EGNN) | 0.10 |
| Feature weight ($\lambda_{\text{feat}}$, EGNN) | 0.90 |
| Structural weight ($\lambda_{\text{coord}}$, ESM-2) | 0.15 |
| Feature weight ($\lambda_{\text{feat}}$, ESM-2) | 0.85 |
| Structural weight ($\lambda_{\text{coord}}$, BLOSUM) | 0.90 |
| Feature weight ($\lambda_{\text{feat}}$, BLOSUM) | 0.10 |

## G Lambda Parameter in Loss Normalization

For coordinate-based losses, *λ* was chosen to reflect the expected spatial separation between unrelated residues. Empirically, distances between unrelated *C*_*α*_ atoms typically fall within the range of approximately 10-30 Å[51]. We therefore set the default value to *λ* = 20 Å, corresponding to the average distance within this range. To ensure that this choice does not artificially influence the evaluation of TM-align, we assessed the robustness of the normalization across a range of penalty values spanning *λ* ∈ [10, 30] Å. For each value, normalized coordinate losses were recomputed and the resulting TM-align rankings were compared. Rank-based summary statistics demonstrate that the relative ordering of TM-align alignments remains stable across this range (Table G1), indicating that results are not overly sensitive to the specific choice of *λ*.

For the BLOSUM loss *λ* was defined analogously as a fixed penalty for unmatched residues, but with a different interpretation. In this case, *λ* corresponds to the maximum possible substitution loss in the BLOSUM62 matrix. Using the highest observed substitution score (+11) and the most negative mismatch score (− 4), which reflects amino acid substitutions that are strongly disfavored and unlikely to occur in nature relative to chance [52], the maximum per-residue loss was computed as

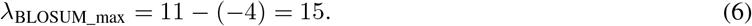

This choice ensures that uncovered residues contribute a worst-case substitution loss consistent with the empirical evolutionary constraints encoded in the scoring matrix.

**Table G1:** Stability of TM-align coordinate-loss rankings across unaligned-residue penalties. Rank correlation of TM-align normalized coordinate losses across different values of the unaligned-residue penalty *λ*, computed relative to the reference value *λ* = 20 Å. Spearman’s *ρ* and Kendall’s *τ* quantify the stability of TM-align loss rankings with respect to changes in *λ*, indicating that relative ordering of TM-align alignments is robust across a broad range of penalty values.

| Penalty $\lambda$ (Å) | Spearman $\rho$ | Kendall $\tau$ |
| --- | --- | --- |
| 10 | 0.999 | 0.969 |
| 12 | 0.999 | 0.975 |
| 14 | 0.999 | 0.981 |
| 16 | 1.000 | 0.987 |
| 18 | 1.000 | 0.994 |
| 20 | 1.000 | 1.000 |
| 22 | 1.000 | 0.994 |
| 24 | 1.000 | 0.987 |
| 26 | 0.999 | 0.981 |
| 28 | 0.999 | 0.974 |
| 30 | 0.998 | 0.968 |

### G.1 Clustering of EpiRanha Predictions

To assess whether EpiRanha identifies multiple distinct epitope-like regions on the surface of a given antigen, we clustered independent epitope predictions generated for the same epitope–antigen pair. Each predicted epitope was represented as a set of antigen residue indices corresponding to a single EpiRanha run. Pairwise dissimilarities between epitope predictions were computed using the Jaccard distance, defined as one minus the ratio of the intersection to the union of residue sets. For each epitope–antigen pair, we constructed a precomputed distance matrix and applied agglomerative hierarchical clustering with average linkage and a fixed distance threshold, allowing the number of clusters to be determined automatically.

Cluster quality was assessed using the silhouette score and by comparing mean intra-and inter-cluster distances. For each cluster, a representative epitope was selected as the medoid, defined as the cluster member with minimal mean distance to all other members. This procedure enabled us to determine whether repeated EpiRanha runs converged on a single epitope-like region or consistently identified multiple distinct regions on the antigen surface.

To quantify the spatial separation between distinct epitope-like regions identified for the same antigen, we computed geometric distances between cluster representatives. For epitope–antigen pairs yielding exactly two clusters, each cluster was represented by its medoid epitope. Using the corresponding antigen structure, we extracted C_*α*_ coordinates for all residues in each medoid and computed the Euclidean distance between the centroids of the two residue sets. This centroid-to-centroid distance provides a coarse measure of the spatial separation between epitope-like regions on the antigen surface. As an additional sanity check, we also computed the minimum C_*α*_-C_*α*_ distance between any residue pair across the two medoids.

## H Experimental Setup

All experiments in this study were conducted in Conda virtual environments to ensure reproducibility and library dependency management. The EGNN model was adapted from an existing PyTorch EGNN implementation [53], with modifications tailored to EpiRanha. Key libraries included PyTorch, BioPython, NumPy, Joblib, heapq, Optuna, scikit-learn, SciPy, Matplotlib, and Pandas. Moreover, datasets were stored in Amazon S3 buckets for efficient access during training and evaluation. Weights & Biases was used to track models, while GitLab was utilized for proper version control. All source code is available in the corresponding GitLab repository.

TM-align was analysed on an AWS EC2 m5.large instance featuring 2 vCPUs. The EGNN hyperparameter search was performed on an AWS EC2 g4dn.xlarge instance, which provides a single NVIDIA T4 GPU. A faster instance or better parallelization could have reduced runtime. For model training, a Slurm cluster with multiple g5.2xlarge instances, each equipped with an NVIDIA A10G GPU, was used to handle the computational load efficiently. Training all eight model variants with cross-validation took approximately 20 hours using 20 parallel instances. Epitope matching with EpiRanha was distributed across 192 vCPUs of a c7a.48xlarge instance to allow large-scale parallelization.

## I Heatmaps Data Curation and Filtering

For the all-vs-all heatmap analysis, a small subset of antigens was excluded due to insufficient antigen length relative to the epitope size, resulting in a large fraction of missing alignments. Specifically, antigens `8pii_B`, `6s0y_C`, `8dtn_B`, and `1mvf_D` were omitted, as more than 70% (or 20% for `1mvf_D`) of their epitope-antigen comparisons were undefined. To preserve the interpretability of cognate (self-matching) cells in the heatmaps, the corresponding epitopes were also removed from the analysis. These cases were retained for self-matching and runtime analyses, where the native epitope–antigen comparison is always well-defined.

## J TM-align Failure Modes: Influence of Epitope and Antigen Length

To better understand the factors underlying TM-align failures on cognate epitope-antigen pairs recovery beyond epitope discontiguity alone, we examined the influence of epitope and antigen length on alignment loss. As shown in Figure J1a, epitope length does not appear to explain why some highly discontiguous epitopes exhibit near-zero TM-align loss while others incur substantial loss. Discontiguity spans a similar range across short and long epitopes, and elevated loss is observed only in a subset of cases, indicating that epitope size alone is not a determining factor. Antigen length (Figure J1b) likewise shows no strong global correlation with loss; however, structural inspection of representative cases (Figure J2) suggests that TM-align tends to struggle more as the effective search space increases. In larger, multi-domain antigens, the broader structural context may introduce additional competing alignments, increasing the likelihood of suboptimal global superpositions and consequently higher loss.

**Figure J1:**
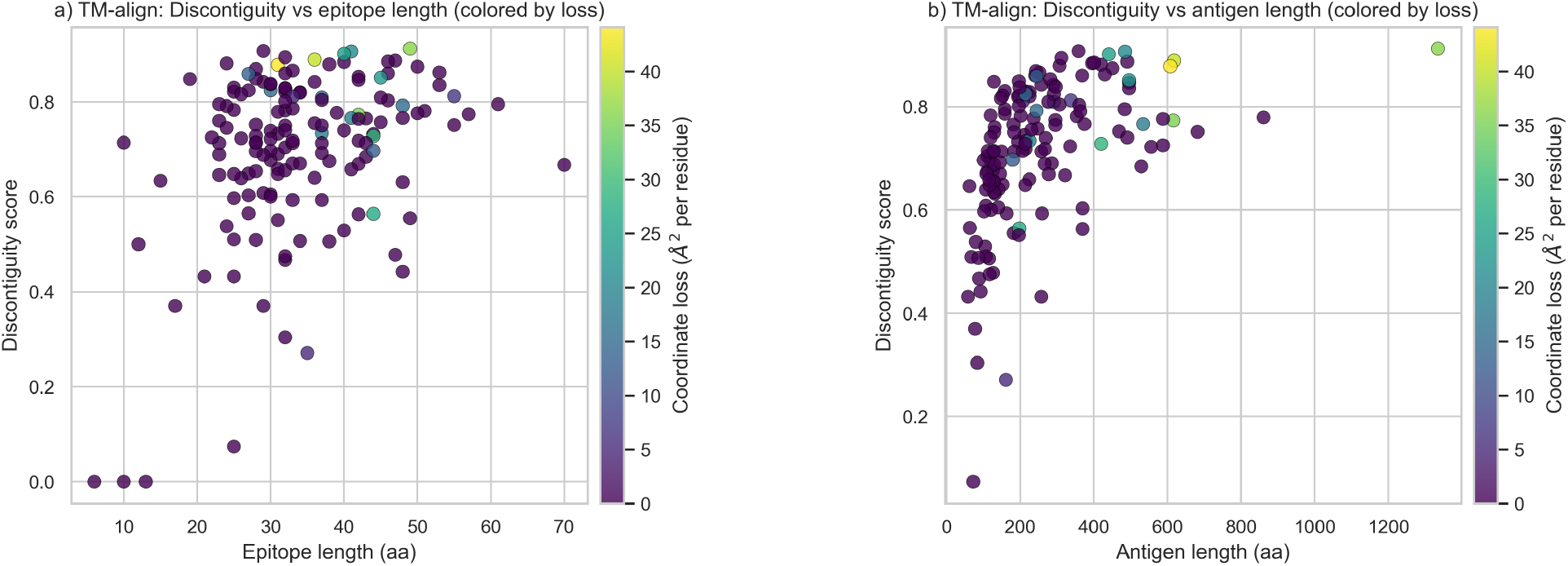
Relationship between epitope length, antigen length, epitope discontiguity, and TM-align loss. Each point represents an epitope–antigen alignment evaluated using TM-align. The y-axis denotes the epitope structural discontiguity score. In (a), the x-axis corresponds to epitope length (amino acids), and in (b), to antigen length (amino acids). Color encodes coordinate loss (Å^2^ per residue).

**Figure J2:**
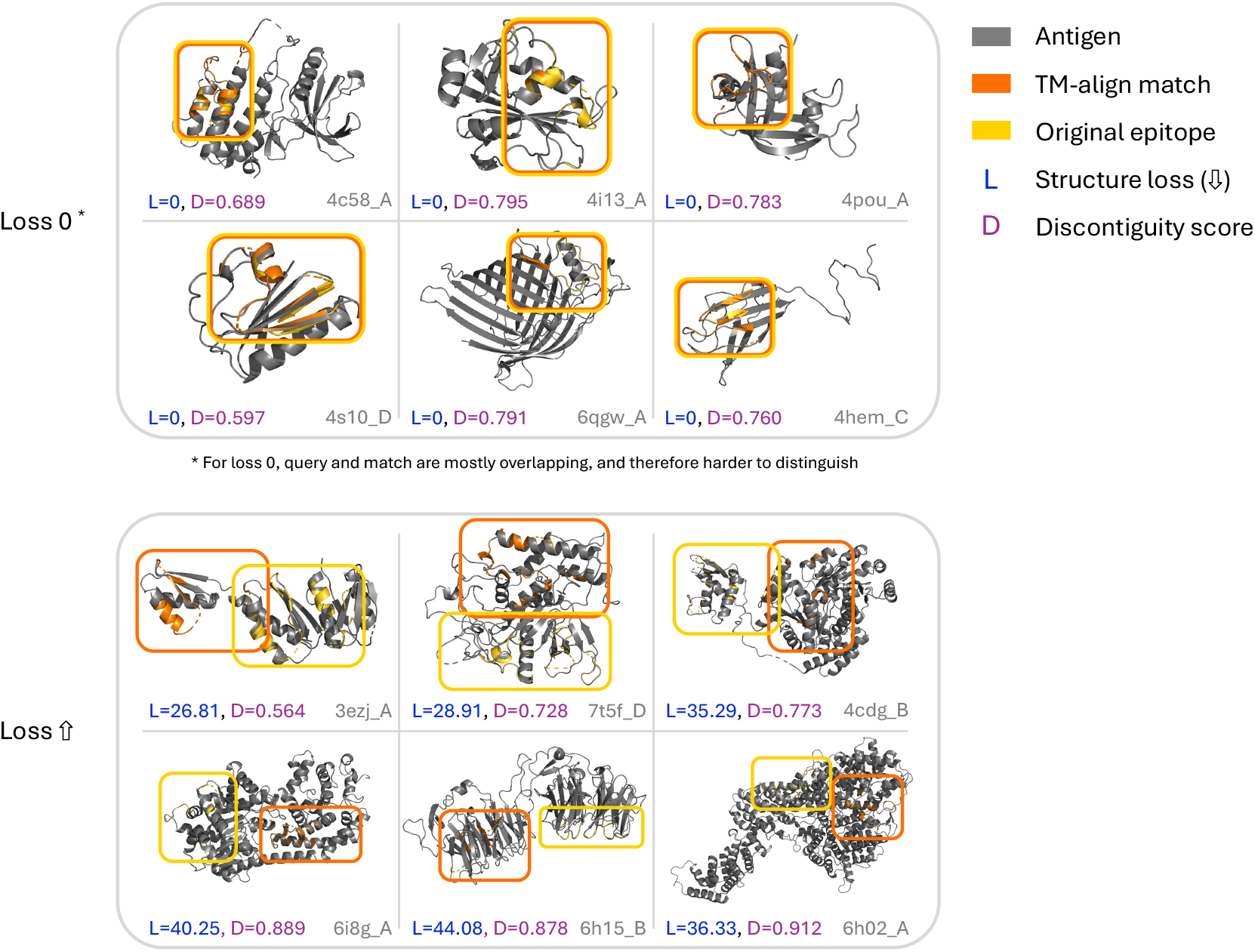
Structural visualization of representative examples illustrating the relationship between TM-align loss and epitope discontiguity. Representative epitope–antigen alignments illustrating the relationship between structural loss (L) and epitope discontiguity (D). The upper examples show cases where TM-align reports near-zero structural loss (*L* = 0) despite high epitope discontiguity. The lower examples exhibit both high structural loss (*L >* 25 Å^2^) and high discontiguity.

**Figure K1:**
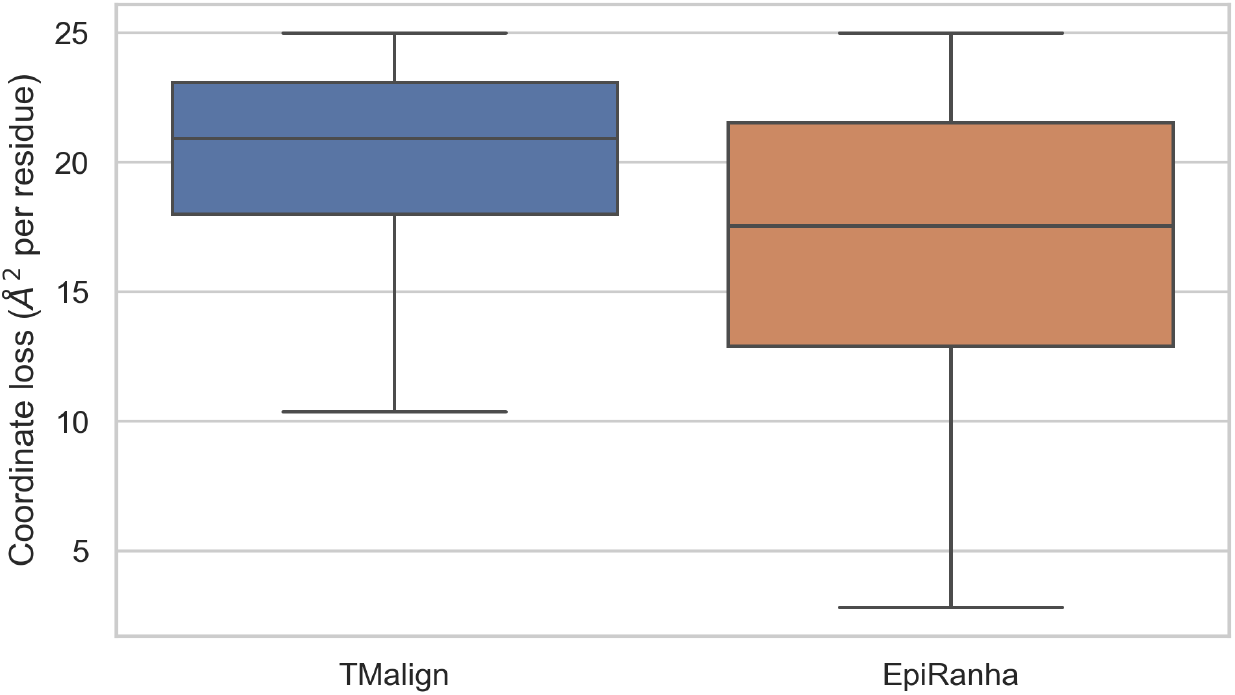
Distribution of cross-antigen alignment losses below the 25 Å threshold. Coordinate loss (Å^2^ per residue) for non-cognate epitope–antigen pairs identified by TM-align and EpiRanha with losses below 25 Å. Each box summarizes the distribution across all cross-antigen matches meeting this criterion.

## K Analysis of Cross-Antigen Matches

To complement these representative examples, we quantified the number of non-self epitope–antigen pairs identified by each method with coordinate losses below the 25 Å threshold (Figure K1). This cutoff reflects structurally meaningful matches under our evaluation framework. The resulting distribution highlights systematic differences in cross-antigen matching behavior between the two approaches. While both methods recover structurally similar regions across non-cognate antigens, EpiRanha yields a broader distribution of low-loss matches, consistent with its flexible residue-level correspondence strategy. In contrast, TM-align’s rigid global superposition constrains its ability to identify certain epitope-scale similarities.

### K.1 Qualitative Analysis of Representative Cross-Antigen Matches

While the main results focus on quantitative trends across all epitope-antigen pairs, off-diagonal entries in the similarity heatmaps can be further examined through representative qualitative examples. These alignments do not correspond to known binding events and are therefore interpreted as indicative of structural similarity rather than recovery of ground-truth epitopes.

Figure 6 shows three representative cross-antigen matching cases. In the first case, the query epitope is highly discontinuous and composed primarily of *β*-strand elements, while the corresponding region identified by TM-align on the search antigen consists predominantly of an *α*-helical segment. TM-align attempts to align these incompatible secondary structure elements, resulting in a high coordinate loss. In contrast, EpiRanha identifies a spatially coherent surface patch that more closely matches the epitope geometry.

The second case illustrates a scenario in which TM-align achieves a lower coordinate loss than EpiRanha. Here, the query epitope is largely coil-rich, and the alignment identified by TM-align involves flexible regions with limited secondary structure. Such regions can be ambiguous to interpret structurally, and although TM-align attains a lower loss in this case, the resulting value remains relatively high.

In the third case, both methods identify cross-antigen matches with coordinate losses below the 25 Å cutoff. However, EpiRanha achieves a substantially lower loss than TM-align. Visual inspection suggests that TM-align aligns an *α*-helical segment of the query epitope to *β*-sheet regions of the target *β*-barrel, whereas EpiRanha recovers a residue correspondence that better preserves local structural context.

Together, these examples illustrate the range of behaviors observed in cross-antigen matching. While TM-align can recover low-loss alignments in specific cases, its matches are constrained by rigid backbone superposition and sensitivity to secondary structure context. EpiRanha more frequently identifies epitope-scale surface patches across non-cognate antigens, reflecting its flexible, residue-level matching strategy.

**Figure L1:**
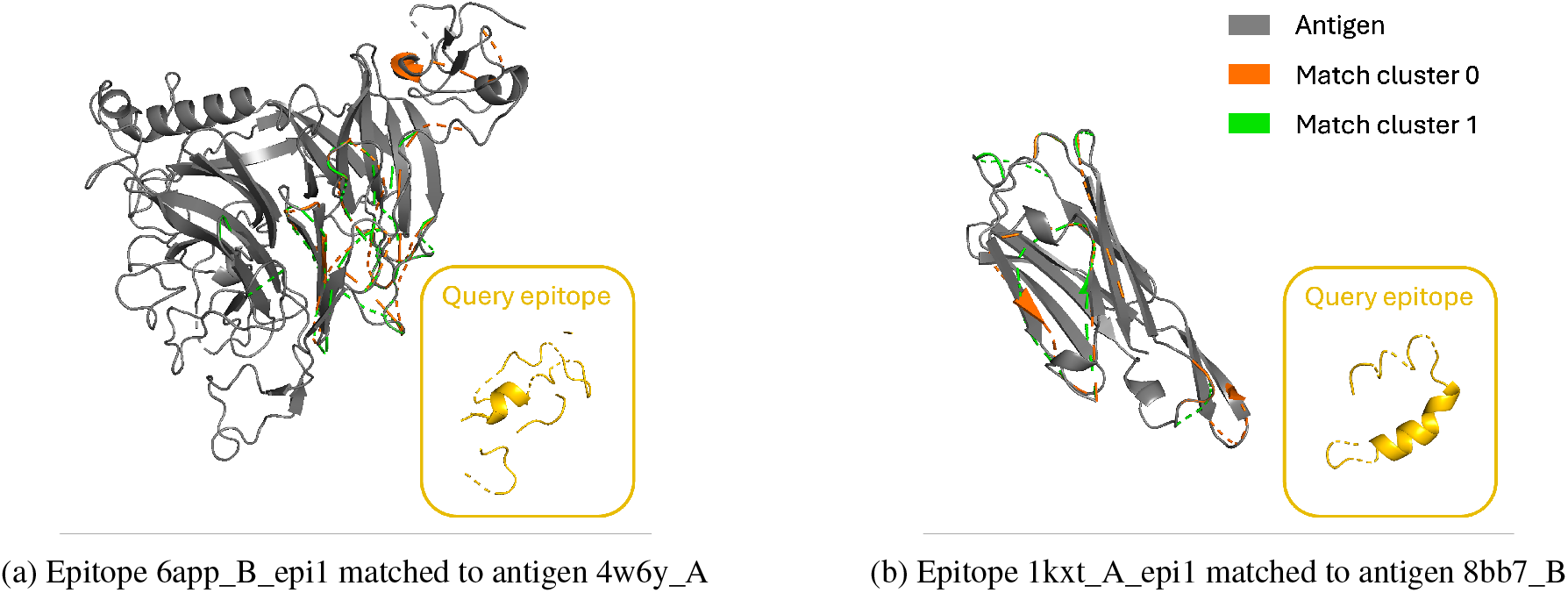
Examples of multiple epitope-like matches identified on a single antigen by EpiRanha. For each epitope-antigen pair, two distinct high-quality matches are shown, corresponding to alternative surface regions on the same antigen. The matches differ in spatial location while maintaining comparable geometric and biochemical consistency, illustrating EpiRanha’s ability to recover multiple plausible epitope-scale regions within a single antigen structure.

## L Multiple Epitope-Like Matches on a Single Antigen

In addition to identifying a single best match for each epitope–antigen pair, EpiRanha frequently detects multiple distinct epitope-like regions on the same antigen surface. This behavior arises naturally from its residue-level scoring and flexible matching strategy, which does not enforce a single global alignment. TM-align, by contrast, is inherently limited to producing a single alignment per epitope–antigen pair, as it optimizes a single rigid-body superposition.

Quantitative support for multiple distinct epitope-like matches on a single antigen is summarized in Table L1. The reported examples exhibit well-separated alignment clusters (silhouette scores *>* 0.4) with limited coordinate loss (*<* 25 Å), indicating that alternative matches are both statistically supported and geometrically plausible. To further assess whether these matches correspond to distinct surface regions, we report centroid–centroid distances between cluster medoids in antigen C*α* coordinate space, providing an interpretable measure of spatial separation without altering the matching procedure. Across representative cases, centroid distances range from near-overlapping patches to clearly distinct regions, consistent with visualizations shown in Figure L1. These results are intended as a capability demonstration rather than evidence of biological ground truth.

Taken together, this analysis highlights that EpiRanha’s matching formulation can yield multiple well-supported and spatially separated solutions for a single epitope–antigen pair, underscoring the diversity of epitope-scale surface correspondences that may be recovered without imposing a single global alignment.

**Table L1:** Clustering and geometric separation statistics for representative multiple-match cases. Each epi-tope–antigen pair yields two alternative alignment clusters identified by EpiRanha. The silhouette score quantifies the quality of the two-cluster partition in alignment space. The centroid distance reports the Euclidean distance between cluster medoids in antigen C*α* coordinate space, serving as an interpretable measure of spatial separation between surface patches. Normalized coordinate loss (in Å per residue) reflects the geometric plausibility of each cluster representative.

| Epitope ID | Antigen ID | Cluster | Silhouette | Centroid dist. (Å) | Coord. loss (Å/res) |
| --- | --- | --- | --- | --- | --- |
| 7t5f_D_epi2 | 1kxt_A | 0 | 0.666 | 4.837 | 20.651 |
| 7t5f_D_epi2 | 1kxt_A | 1 | 0.666 | 4.837 | 20.705 |
| 6app_B_epi1 | 4w6y_A | 0 | 0.502 | 9.797 | 11.813 |
| 6app_B_epi1 | 4w6y_A | 1 | 0.502 | 9.797 | 14.181 |
| 1kxt_A_epi1 | 8bb7_B | 0 | 0.496 | 5.802 | 13.734 |
| 1kxt_A_epi1 | 8bb7_B | 1 | 0.496 | 5.802 | 12.599 |
| 6r7t_B_epi1 | 8bf4_F | 0 | 0.484 | 1.768 | 10.370 |
| 6r7t_B_epi1 | 8bf4_F | 1 | 0.484 | 1.768 | 10.891 |

## M Supplementary Methods

### M.1 Dataset scale

**Figure M1:**
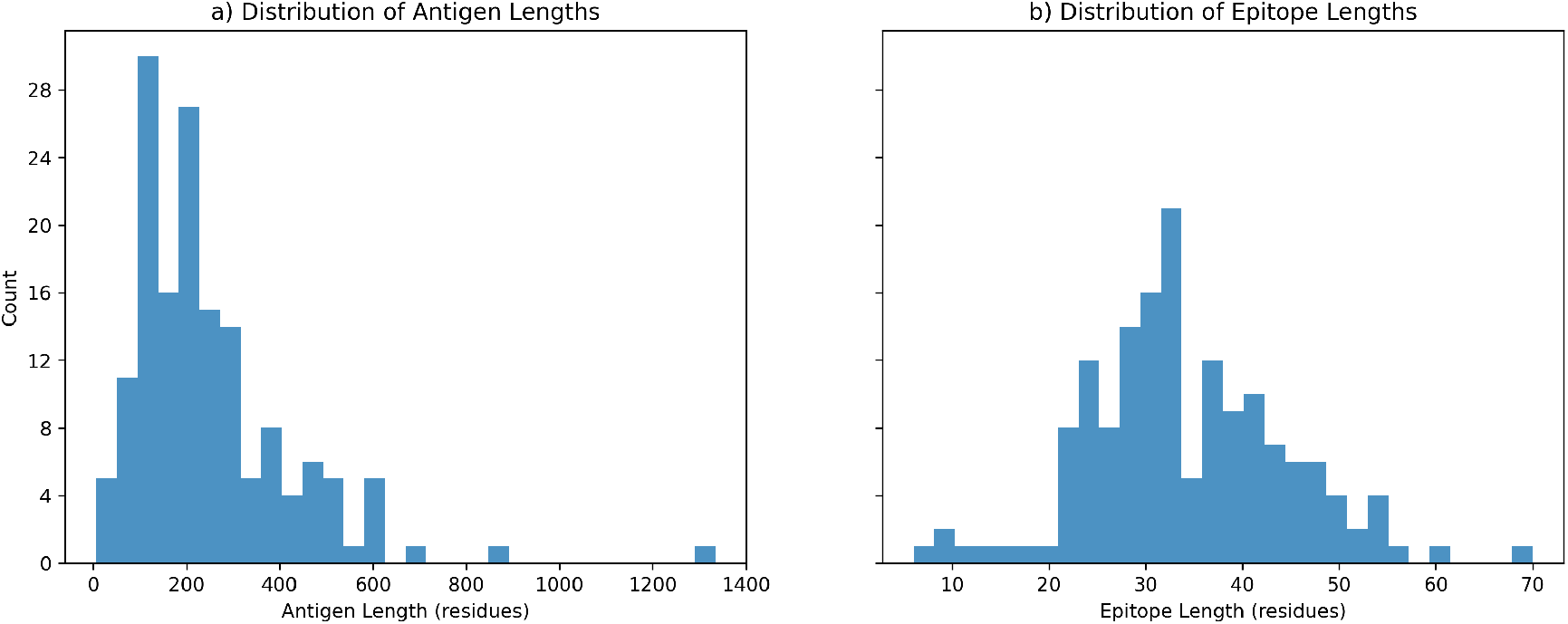
Distribution of antigen and epitope sizes in the SabDab-Nano dataset. Histograms show the number of residues per antigen (left) and per epitope (right) across all analyzed SabDab-Nano complexes. These distributions illustrate the broad range of antigen sizes and the typical scale of epitope regions evaluated in this study.

### M.2 Epitope discontiguity metric

**Figure M2:**
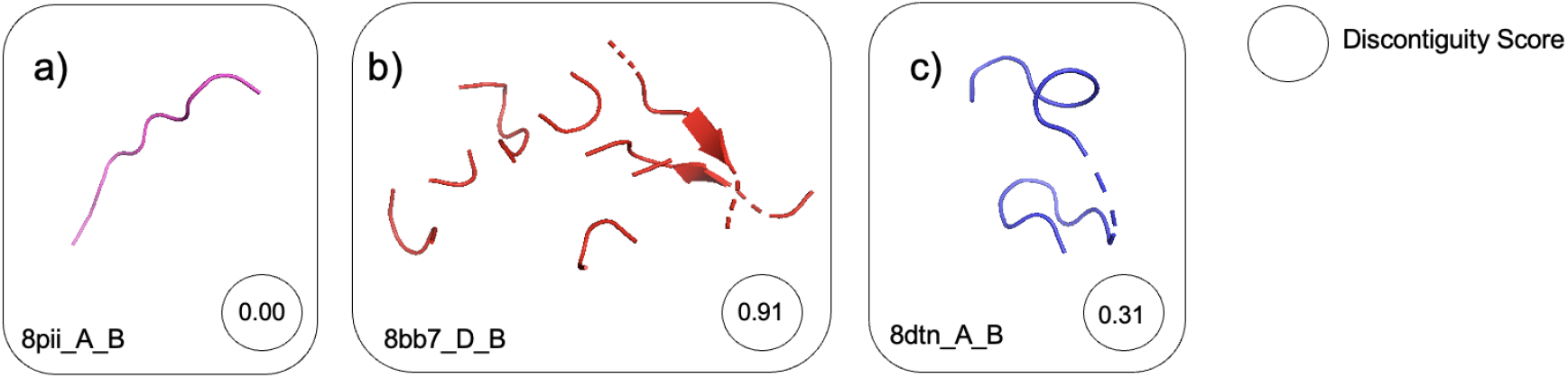
Representative examples illustrating the behavior of the epitope discontiguity metric. Three epitopes from antibody–antigen complexes are shown with increasing degrees of spatial fragmentation (i.e., discontiguity). Panel (a) illustrates a contiguous epitope, (b) a highly fragmented conformational epitope, and (c) a moderately discontiguous case. The discontiguity score (shown in each panel) increases with the degree of epitope fragmentation.

### M.3 Index-Epitope and Index-Antigen Mapping

**Table M1:**
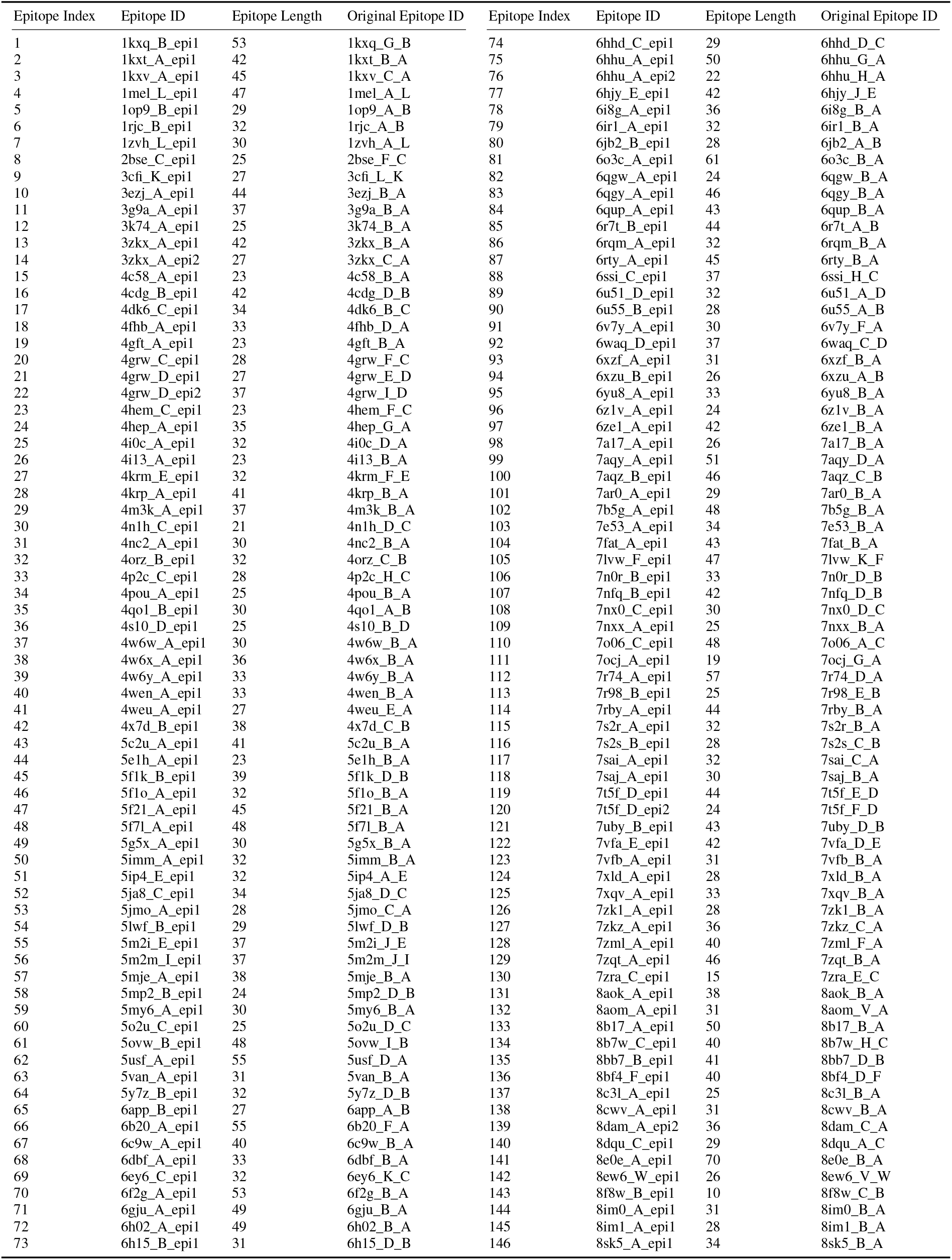
Epitope index mapping between benchmark epitope identifiers and original PDB IDs (twin-column presentation)

**Table M2:**
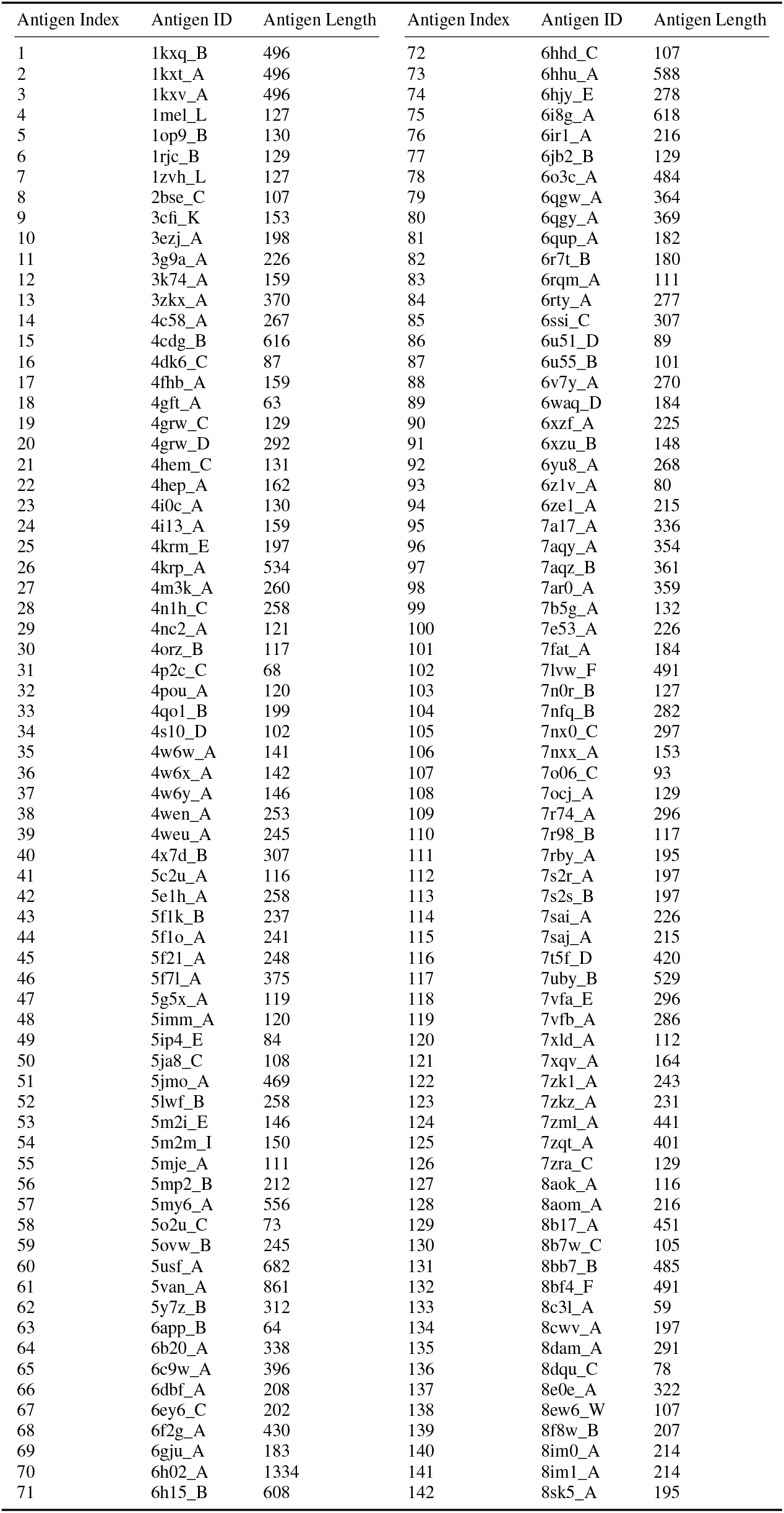
Antigen index mapping (twin-column presentation)

## Notes

### Competing Interest Statement

The authors have declared no competing interest.

## References

[1] Ruei-Min Lu, Yu-Chyi Hwang, I-Ju Liu, Chi-Chiu Lee, Han-Zen Tsai, Hsin-Jung Li, and Han-Chung Wu. Development of therapeutic antibodies for the treatment of diseases. Journal of biomedical science, 27:1–30, 2020.

[2] María Sofía Castelli, Paul McGonigle, and Pamela J Hornby. The pharmacology and therapeutic applications of monoclonal antibodies. Pharmacology research & perspectives, 7(6):e00535, 2019.

[3] Md Harun-Or-Rashid, Most Nazmin Aktar, Md Sabbir Hossain, Nadia Sarkar, Md Rezaul Islam, Md Easin Arafat, Shukanta Bhowmik, and Shin-ichi Yusa. Recent advances in micro-and nano-drug delivery systems based on natural and synthetic biomaterials. Polymers, 15(23):4563, 2023.

[4] Nimrod D. Rubinstein, Itay Mayrose, Dan Halperin, Daniel Yekutieli, Jonathan M. Gershoni, and Tal Pupko. Computational characterization of b-cell epitopes. Molecular Immunology, 45(12):3477–3489, 7 2008.

[5] Neil S Greenspan and Enrico Di Cera. Defining epitopes: It’s not as easy as it seems. Nature Biotechnology, 17(10):936–937, 1999.

[6] Astrid Musnier, Thomas Bourquard, Amandine Vallet, Laetitia Mathias, Gilles Bruneau, Mohammed Akli Ayoub, Ophélie Travert, Yannick Corde, Nathalie Gallay, Thomas Boulo, et al. A new in silico antibody similarity measure both identifies large sets of epitope binders with distinct cdrs and accurately predicts off-target reactivity. International journal of molecular sciences, 23(17):9765, 2022.

[7] Andre F Fonseca and Dinler A Antunes. Crossdome: an interactive r package to predict cross-reactivity risk using immunopeptidomics databases. Frontiers in Immunology, 14:1142573, 2023.

[8] Salvador Eugenio C Caoili. Expressing redundancy among linear-epitope sequence data based on residue-level physicochemical similarity in the context of antigenic cross-reaction. Advances in Bioinformatics, 2016(1):1276594, 2016.

[9] Fatima Ferreira, Thomas Hawranek, P Gruber, N Wopfner, and A Mari. Allergic cross-reactivity: from gene to the clinic. Allergy, 59(3):243–267, 2004.

[10] John Ingraham, Vikas Garg, Regina Barzilay, and Tommi Jaakkola. Generative models for graph-based protein design. In H. Wallach, H. Larochelle, A. Beygelzimer, F. d’Alché-Buc, E. Fox, and R. Garnett, editors, Advances in Neural Information Processing Systems, volume 32. Curran Associates, Inc., 2019.

[11] Justas Dauparas, Ivan Anishchenko, Nathaniel Bennett, Hua Bai, Robert J Ragotte, Lukas F Milles, Basile IM Wicky, Alexis Courbet, Rob J de Haas, Neville Bethel, et al. Robust deep learning–based protein sequence design using proteinmpnn. Science, 378(6615):49–56, 2022.

[12] Ovidiu Ivanciuc, Catherine H Schein, Tzintzuni Garcia, Numan Oezguen, Surendra S Negi, and Werner Braun. Structural analysis of linear and conformational epitopes of allergens. Regulatory toxicology and pharmacology, 54(3):S11–S19, 2009.

[13] Thiruvarangan Ramaraj, Thomas Angel, Edward A Dratz, Algirdas J Jesaitis, and Brendan Mumey. Antigen– antibody interface properties: Composition, residue interactions, and features of 53 non-redundant structures. Biochimica et Biophysica Acta (BBA)-Proteins and Proteomics, 1824(3):520–532, 2012.

[14] Pinak Chakrabarti and Joel Janin. Dissecting protein–protein recognition sites. Proteins: Structure, Function, and Bioinformatics, 47(3):334–343, 2002.

[15] Yang Zhang and Jeffrey Skolnick. Scoring function for automated assessment of protein structure template quality. Proteins: Structure, Function, and Bioinformatics, 57(4):702–710, 2004.

[16] Mindaugas Margelevičius. Gtalign: Spatial index-driven protein structure alignment, superposition, and search. Nature Communications, 15(1):7305, 2024.

[17] Michel van Kempen, Stephanie S. Kim, Charlotte Tumescheit, Milot Mirdita, Jeongjae Lee, Cameron L.M. Gilchrist, Johannes Söding, and Martin Steinegger. Fast and accurate protein structure search with foldseek. Nature Biotechnology, 42(2):243–246, 2 2024.

[18] Yang Zhang and Jeffrey Skolnick. Tm-align: A protein structure alignment algorithm based on the tm-score. Nucleic Acids Research, 33(7):2302–2309, 2005.

[19] Shashi Bhushan Pandit and Jeffrey Skolnick. Fr-tm-align: a new protein structural alignment method based on fragment alignments and the tm-score. BMC bioinformatics, 9:1–11, 2008.

[20] Hyunbin Kim, Rachel Seongeun Kim, Milot Mirdita, and Martin Steinegger. Structural motif search across the protein-universe with folddisco. bioRxiv, pages 2025–07, 2025.

[21] Tomonori Osajima, Masaaki Suzuki, Saburo Neya, and Tyuji Hoshino. Computational and statistical study on the molecular interaction between antigen and antibody. Journal of Molecular Graphics and Modelling, 53:128–139, 2014.

[22] P. Gainza, F. Sverrisson, F. Monti, E. Rodolà, D. Boscaini, M. M. Bronstein, and B. E. Correia. Deciphering interaction fingerprints from protein molecular surfaces using geometric deep learning. Nature Methods, 17(2):184–192, 2 2020.

[23] Freyr Sverrisson, Jean Feydy, Bruno E Correia, and Michael M Bronstein. Fast end-to-end learning on protein surfaces. In Proceedings of the IEEE/CVF Conference on Computer Vision and Pattern Recognition, pages 15272–15281, 2021.

[24] Zeming Lin, Halil Akin, Roshan Rao, Brian Hie, Zhongkai Zhu, Wenting Lu, Allan dos Santos Costa, Maryam Fazel-Zarandi, Tom Sercu, Sal Candido, et al. Language models of protein sequences at the scale of evolution enable accurate structure prediction. BioRxiv, 2022:500902, 2022.

[25] Yuansong Zeng, Zhuoyi Wei, Qianmu Yuan, Sheng Chen, Weijiang Yu, Yutong Lu, Jianzhao Gao, and Yue-dong Yang. Identifying b-cell epitopes using alphafold2 predicted structures and pretrained language model. Bioinformatics, 39(4):btad187, 4 2023.

[26] Constantin Schneider, Matthew I.J. Raybould, and Charlotte M. Deane. Sabdab in the age of biotherapeutics: Updates including sabdab-nano, the nanobody structure tracker. Nucleic Acids Research, 50, 2022.

[27] Helen M Berman, John Westbrook, Zukang Feng, Gary Gilliland, Talapady N Bhat, Helge Weissig, Ilya N Shindyalov, and Philip E Bourne. The protein data bank. Nucleic acids research, 28(1):235–242, 2000.

[28] John Jumper, Richard Evans, Alexander Pritzel, Tim Green, Michael Figurnov, Olaf Ronneberger, Kathryn Tunyasuvunakool, Russ Bates, Augustin Žídek, Anna Potapenko, et al. Highly accurate protein structure prediction with alphafold. nature, 596(7873):583–589, 2021.

[29] Zeming Lin, Halil Akin, Roshan Rao, Brian Hie, Zhongkai Zhu, Wenting Lu, Nikita Smetanin, Robert Verkuil, Ori Kabeli, Yaniv Shmueli, et al. Evolutionary-scale prediction of atomic-level protein structure with a language model. Science, 379(6637):1123–1130, 2023.

[30] Victor Garcia Satorras, Emiel Hoogeboom, and Max Welling. E(n) equivariant graph neural networks. In Proceedings of the 38th International Conference on Machine Learning, volume 139 of Proceedings of Machine Learning Research, pages 9323–9332. PMLR, 2021.

[31] David F Crouse. On implementing 2d rectangular assignment algorithms. IEEE Transactions on Aerospace and Electronic Systems, 52(4):1679–1696, 2016.

[32] Osamu Gotoh. An improved algorithm for matching biological sequences. Journal of molecular biology, 162(3):705–708, 1982.

[33] Magnus Haraldson Høie, Frederik Steensgaard Gade, Julie Maria Johansen, Charlotte Würtzen, Ole Winther, Morten Nielsen, and Paolo Marcatili. Discotope-3.0: improved b-cell epitope prediction using inverse folding latent representations. Frontiers in Immunology, 15:1322712, 2024.

[34] Joakim Nøddeskov Clifford, Magnus Haraldson Høie, Sebastian Deleuran, Bjoern Peters, Morten Nielsen, and Paolo Marcatili. Bepipred-3.0: Improved b-cell epitope prediction using protein language models. Protein Science, 31(12):e4497, 2022.

[35] Veronika I Zarnitsyna, Jennie Lavine, Ali Ellebedy, Rafi Ahmed, and Rustom Antia. Multi-epitope models explain how pre-existing antibodies affect the generation of broadly protective responses to influenza. PLoS pathogens, 12(6):e1005692, 2016.

[36] Inbal Sela-Culang, Vered Kunik, and Yanay Ofran. The structural basis of antibody-antigen recognition. Frontiers in immunology, 4:302, 2013.

[37] Saleh Riahi, Jae Hyeon Lee, Taylor Sorenson, Shuai Wei, Sven Jager, Reza Olfati-Saber, Yanfeng Zhou, Anna Park, Maria Wendt, Herve Minoux, and Yu Qiu. Surface id: A geometry-aware system for protein molecular surface comparison. Bioinformatics, 39(4):btad196, 4 2023.

[38] Woosub Kim, Milot Mirdita, Eli Levy Karin, Cameron LM Gilchrist, Hugo Schweke, Johannes Söding, Emmanuel D Levy, and Martin Steinegger. Rapid and sensitive protein complex alignment with foldseek-multimer. Nature Methods, pages 1–4, 2025.

[39] Pablo Gainza, Sarah Wehrle, Alexandra Van Hall-Beauvais, Anthony Marchand, Andreas Scheck, Zander Harteveld, Stephen Buckley, Dongchun Ni, Shuguang Tan, Freyr Sverrisson, et al. De novo design of protein interactions with learned surface fingerprints. Nature, 617(7959):176–184, 2023.

[40] Qianmu Yuan, Jianwen Chen, Huiying Zhao, Yaoqi Zhou, and Yuedong Yang. Structure-aware protein-protein interaction site prediction using deep graph convolutional network. Bioinformatics, 38:125–132, 1 2022.

[41] Manon Réau, Nicolas Renaud, Li C. Xue, and Alexandre M.J.J. Bonvin. Deeprank-gnn: a graph neural network framework to learn patterns in protein–protein interfaces. Bioinformatics, 39, 1 2023.

[42] Alexey Strokach, David Becerra, Carles Corbi-Verge, Albert Perez-Riba, and Philip M. Kim. Fast and flexible protein design using deep graph neural networks. Cell Systems, 11:402–411.e4, 10 2020.

[43] Wenhao Gao, Sai Pooja Mahajan, Jeremias Sulam, and Jeffrey J. Gray. Deep learning in protein structural modeling and design, 12 2020.

[44] Fabian Traxler. Antibody-Antigen Binding Affinity Prediction through the use of geometric deep learning. PhD thesis, Wien, 2023.

[45] Joe G Greener and Kiarash Jamali. Fast protein structure searching using structure graph embeddings. Bioinformatics Advances, 5(1):vbaf042, 2025.

[46] Lasse M Blaabjerg, Nicolas Jonsson, Wouter Boomsma, Amelie Stein, and Kresten Lindorff-Larsen. A joint embedding of protein sequence and structure enables robust variant effect predictions. bioRxiv, pages 2023–12, 2023.

[47] Joakim Nøddeskov Clifford, Eve Richardson, Bjoern Peters, and Morten Nielsen. Abepitope-1.0: Improved antibody target prediction by use of alphafold and inverse folding. Science Advances, 11(24):eadu1823, 2025.

[48] Peter Eastman, Jason Swails, John D Chodera, Robert T McGibbon, Yutong Zhao, Kyle A Beauchamp, Lee-Ping Wang, Andrew C Simmonett, Matthew P Harrigan, Chaya D Stern, et al. Openmm 7: Rapid development of high performance algorithms for molecular dynamics. PLoS computational biology, 13(7):e1005659, 2017.

[49] Rita Casadio, Pier Luigi Martelli, and Castrense Savojardo. Machine learning solutions for predicting protein–protein interactions, 11 2022.

[50] Takuya Akiba, Shotaro Sano, Toshihiko Yanase, Takeru Ohta, and Masanori Koyama. Optuna: A next-generation hyperparameter optimization framework. In Proceedings of the 25th ACM SIGKDD International Conference on Knowledge Discovery and Data Mining, 2019.

[51] Jianyi Yang, Ivan Anishchenko, Hahnbeom Park, Zhenling Peng, Sergey Ovchinnikov, and David Baker. Improved protein structure prediction using predicted interresidue orientations. Proceedings of the National Academy of Sciences, 117(3):1496–1503, 2020.

[52] Steven Henikoff and Jorja G Henikoff. Amino acid substitution matrices from protein blocks. Proceedings of the National Academy of Sciences, 89(22):10915–10919, 1992.

[53] Phil Wang. Egnn - pytorch: E(n)-equivariant graph neural networks in pytorch. https://github.com/lucidrains/egnn-pytorch, 2021. Accessed: September 2024.

